# Systems-level characterization of EGFR kinase inhibitors reveals heterogeneous effects on Mtb-macrophage interactions

**DOI:** 10.1101/2025.10.17.683041

**Authors:** Cal Gunnarsson, Rachel A. McGinn, Jacob Hochfelder, Bryan D. Bryson

**Affiliations:** Department of Biological Engineering, Massachusetts Institute of Technology, Cambridge, MA, USA; Ragon Institute of MGH, MIT, and Harvard, Cambridge, MA, USA

## Abstract

*Mycobacterium tuberculosis* (Mtb) causes deadly, antibiotic-recalcitrant disease. Host-directed therapy (HDT) is a proposed, antibiotic-complementary approach that enhances host antimicrobial function, thereby restricting intracellular Mtb. Drug repurposing screens suggest that inhibiting epidermal growth factor receptor (EGFR) may improve macrophage restriction of Mtb. However, the role of EGFR in restriction is not well understood. We show EGFR kinase inhibitors are not generally Mtb-restrictive in human monocyte-derived macrophages. Few EGFR inhibitors restrict intracellular Mtb and do so via heterogeneous mechanisms, including direct antibiotic effects and host-dependent, likely EGFR-independent mechanisms. During host-dependent therapy, intracellular Mtb induce defined, stress-responsive gene modules, with each drug inducing a unique combination of restriction-associated stresses. We decompose intracellular Mtb responses into host-dependent contributions and host-independent contributions suggestive of direct drug-Mtb interaction. Together, our data nuances the host-directed model of EGFR inhibitors for tuberculosis and provides a roadmap for systematically characterizing repurposed drugs from multiple angles of the drug-host-pathogen interaction.

## Introduction

*Mycobacterium tuberculosis* (Mtb) is a deadly and prevalent bacterial pathogen. Infection begins when Mtb are phagocytosed by macrophages. Through subversion of or adaptation to macrophage antimicrobial responses, Mtb persists in the phagosome. Lengthy combination antibiotic therapy, then, is required to clear infection.^1^ Host-directed therapy (HDT) is an emerging treatment approach based on accessing host responses beneficial to the course of disease, such as inducing antimicrobial responses to clear intracellular Mtb, dampening tissue-damaging responses, or improving drug penetration.^2^

One form of large-scale HDT discovery involves repurposing compound libraries, which include host-targeting drugs, and screening on intracellular bacterial growth in drug-treated, infected host cells.^3–5^ Screen hits are hypothesized to be predominantly host-directed if some or all of the following criteria are met: the drug has a known host target, drug treatment changes expression levels or localization of antimicrobial-associated transcripts or proteins, and the drug has no effect on axenic bacterial growth at the same concentration. When known, host targets of screen hits are also hypothesized to be host factors regulating infection, which can be supported by genetic perturbation or use of small molecules with the same target profile. Classifying a drug as a host-directed therapy, then, is a mechanistic claim about where and how intracellular growth restriction arises, and categorization of screen hits as host-directed requires detailed study.

Despite differences in cell line and metric of Mtb restriction, HDT screens have converged on the epidermal growth factor receptor (EGFR) kinase inhibitor gefitinib.^3,5^ Screens have also identified other EGFR kinase inhibitors that restrict intracellular Mtb.^3–5^ However, the mechanism of restriction for gefitinib and for other EGFR inhibitors is not well understood.

The drug-target interaction between EGFR inhibitors and EGFR in human macrophages has not been completely established. EGFR is inconsistently detected in *ex vivo* and *in vitro* macrophages and macrophage-like cells.^6–8^ EGFR inhibitors have emerged from screens on intracellular growth restriction of *Salmonella* and *Staphylococcus aureus*,^9–11^ but in some cases, these effects are EGFR-independent.^11^ Host-targeted screens with human macrophage-like THP-1 cells and *Mycobacterium bovis* Bacillus Calmette-Guerin find no effect of EGFR knockout on macrophage survival during infection.^12^ Establishing whether gefitinib and other EGFR inhibitors proposed as HDTs predominantly act via EGFR would both inform their development as drugs and inform biological knowledge of host factors influencing infection outcome.

Although gefitinib upregulates several antimicrobial-related pathways, not all pathways are required for restriction. Mtb-infected murine macrophages treated with gefitinib show increased STAT3-mediated cytokine signaling and autophagic flux, but knocking out STAT3 or the autophagy effector Atg7 had no effect on gefitinib-mediated restriction.^13^ Functionally, gefitinib treatment increases the number of acidic, proteolytic compartments and enhances killing of Mtb trafficked to these compartments.^13^ However, it’s not known whether acidification directly restricts Mtb or if other pH-dependent processes restrict Mtb. This is a relevant distinction, because Mtb can remain intact and retain viability despite being trafficked to the lysosome.^14,15^ Understanding how HDTs affect bacterial state can inform screening strategies and use in combination with antibiotics.

Using EGFR inhibitors as a prototype, we sought to determine whether HDTs had shared features aside from their effects on intracellular bacterial burden. Despite their shared target, only a subset of EGFR inhibitors restricted intracellular bacteria, and follow-up experiments suggested multiple, off-target mechanisms of action. Several drugs restricted intracellular bacteria by direct antibiotic effects, while others exhibited host-dependent activity. By profiling the intracellular Mtb transcriptional state, we show that growth restriction by host-dependent inhibitors is associated with multiple intracellular bacterial states. These intracellular bacterial states can be decomposed into sets of environment response-associated genes. By profiling axenic Mtb state upon drug treatment, we show that host-dependent inhibitors alter Mtb transcriptional state, despite not driving gross defects in bacterial growth, viability, or metabolic phenotype, and that the intracellular bacterial state can be attributed to both host-independent and host-dependent effects. These data provide a framework for characterizing repurposed compounds and suggest that the host-directed hypothesis for EGFR kinase inhibitors underestimates direct Mtb-drug interactions and overestimates the specific, host-directed role of EGFR kinase inhibition in intracellular growth restriction.

### EGFR inhibitors show different activities against intracellular Mtb

Since multiple independent high-throughput drug repurposing studies identify EGFR kinase inhibitors as hits,^3–5^ we tested whether EGFR inhibition generally promotes restriction of intracellular Mtb within primary human monocyte-derived macrophages (hMDMs). We selected a panel of 18 EGFR inhibitors spanning nanomolar to micromolar equilibrium dissociation constants (K_d_) for wild type EGFR. We hypothesized that at a standard concentration of 10 µM, we would observe a relationship between quantitative features of a drug’s engagement with EGFR and intracellular growth of Mtb. To measure drug effects on intracellular growth of Mtb, we infected hMDMs with luminescent Mtb and tracked luminescence as a proxy for bacterial growth and metabolic activity^16–18^ (Figure 1A). Drugs were added post-phagocytosis to avoid effects on uptake and remained in the media throughout the infection timecourse. At endpoint, we quantified drug effects by scaling luminescence to vehicle, which allowed us to classify drugs as restrictive (scaled luminescence < 1). Of 18 tested EGFR inhibitors, only five – varlitinib, lapatinib, gefitinib, poziotinib, and pelitinib – strongly reduced intracellular luminescence (Figure 1B). Inhibitors with lower K_d_ values, higher selectivities for EGFR, or stronger percent inhibition of EGFR kinase activity were not necessarily more restrictive (Figure 1C; Supplementary Figure 1A). Groups of structurally similar drugs also did not necessarily perform similarly (Figure 1D). Although different modes of EGFR inhibition can drive different host transcriptional responses,^19,20^ we failed to map growth-restrictive EGFR inhibitors into a single host response category (Supplementary Figure 1B). Since features of EGFR inhibition were insufficient to explain intracellular growth restriction, we tested whether human donor-derived macrophages used in our infection model expressed targetable EGFR on their surface. We failed to detect surface expression of EGFR, ErbB2, or ErbB3 above isotype across multiple human donors (Figure 1E), despite detection in A549 cells using the same antibodies. We similarly failed to detect total EGFR or phosphorylation of EGFR in response to EGF stimulation in hMDM cell lysates, despite detecting total EGFR and EGF-specific EGFR phosphorylation in A549 lysates (Supplementary Figure 2). Together, this strongly suggests an off-target mechanism of action.

**Figure 1.**
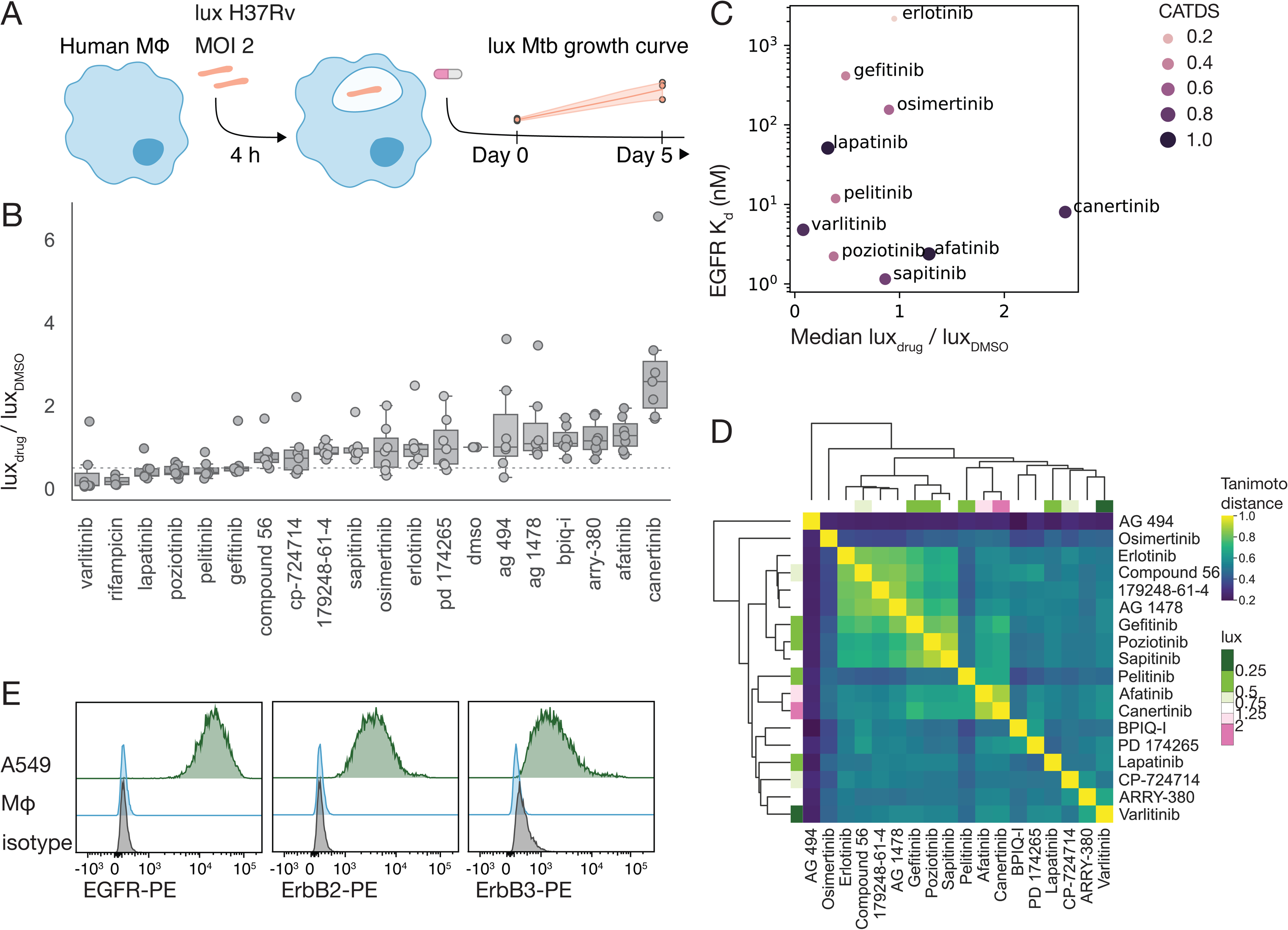
EGFR inhibitors have heterogeneous effects on infection outcome. **A)** Experimental schematic for intracellular Mtb luminescence assays. Human monocyte-derived macrophages were infected with luxABCDE-expressing H37Rv Mtb, treated with drug or vehicle post-phagocytosis, then monitored for luminescence over a 5-day infection. **B)** Intracellular luminescence at day 5, normalized to DMSO for 18 EGFR inhibitors used at 10 µM, plus rifampicin as a positive control. Dotted line corresponds to 50% reduction in intracellular luminescence. Datapoints represent averaged triplicate technical replicates for n=7 donors, 3 independent experiments. **C)** EGFR dissociation constant (K_d_) and selectivity (CATDS) across drugs^92^ is plotted against median effect on intracellular luminescence. **D)** Chemical similarity across drugs. Pairwise Tanimoto similarity is calculated using rdkit, then clustered using Euclidean distance and average linkage. Drugs are annotated according to median effect on intracellular luminescence. **E)** Flow cytometry for surface EGFR, ErbB2, and ErbB3. Representative of n=6 donors.

### Antibiotic effects underlie intracellular restriction by lapatinib and varlitinib

Many EGFR inhibitors are quinazolines, a chemical class that includes several agents optimized for antitubercular activity (Supplementary Figure 3).^21^ Since lapatinib has antitubercular activity,^22^ we tested whether EGFR inhibitors that restricted intracellular growth also restricted axenic growth, using reductive conversion of resazurin in axenic Mtb cultures as a proxy for viable, metabolically active bacteria at endpoint.^23,24^ We cultured Mtb in a microwell format in drug-containing 7H9 broth for 4 days, added resazurin to cultures, and quantified reductive conversion of resazurin after 2 additional days. Since gefitinib and lapatinib are lysosomotropic and could accumulate in the Mtb-containing phagosome,^25^ we tested concentrations up to 50 µM. At concentrations above 10 µM, lapatinib and varlitinib inhibited reductive conversion of resazurin, suggesting direct antitubercular effects, while pelitinib, poziotinib, and gefitinib didn’t inhibit reductive conversion of resazurin at tested concentrations (Figure 2A). To confirm differential drug effects at high concentrations, we monitored growth and intracellular ATP content of Mtb cultures exposed to 25 µM drug (Figure 2B-C). Treatment with lapatinib or varlitinib significantly slowed growth rate (t_1/2_ = 36.7 ± 3.4 h, 25.9 ± 1.6 h; both comparisons adjusted p < 0.0001, ordinary one-way ANOVA), while treatment with pelitinib, poziotinib, and gefitinib slightly slowed growth rate relative to DMSO (t_1/2_ = 18.5h ± 0.5, 20.7 ± 1.5 h, 19.0 ± 0.3 h vs. 16.6 ± 0.7 h DMSO; adjusted p = 0.37, 0.01, 0.21, ordinary one-way ANOVA). This suggests that pelitinib, poziotinib, and gefitinib could alter bacterial viability under conditions not tested here. At 24 hours post-exposure, intracellular ATP content was also significantly lower for cultures treated with varlitinib or lapatinib compared to pelitinib, gefitinib, or poziotinib (Figure 2C). Together, these results distinguish varlitinib and lapatinib as strongly affecting multiple aspects of bacterial state.

**Figure 2.**
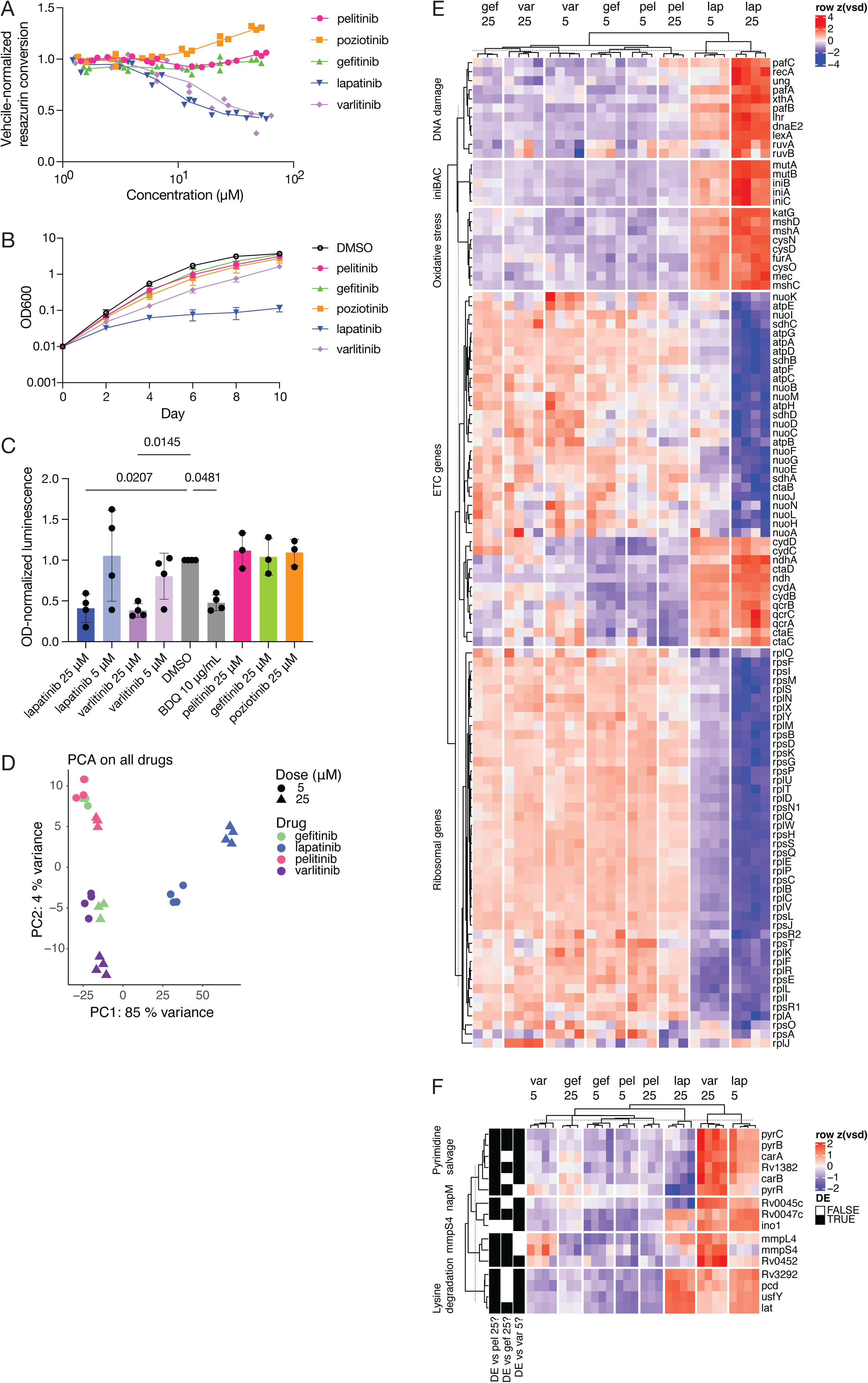
Transcriptional state of lapatinib-treated Mtb supports a directly antibiotic mechanism of intracellular growth restriction. **A)** Effect of drug treatment on axenic Mtb growth and viability, quantified as reductive conversion of resazurin normalized to DMSO. Resazurin was added at 4 days post-treatment, and results were measured via absorbance after 2 additional days. Representative of n=3 independent experiments. **B)** Effect of drug treatment on optical density at a standard concentration of 25 µM. Mean and standard deviation are plotted for n=4 independently generated bacterial stocks measured in 2 independent experiments. **C)** Effect of drug treatment on intracellular ATP content at 24h, measured by BacTiterGlo. Relative luminescence is normalized to input OD, then the DMSO condition. Statistical significance was tested relative to the DMSO control using an ordinary one-way ANOVA and Holm-Šídák’s multiple comparisons test. n=4 independently generated bacterial stocks measured in 2 independent experiments. BDQ = bedaquiline. **D - F)** RNA-seq was performed on n=3-4 independently generated Mtb stocks treated with 5 or 25 µM lapatinib, varlitinib, pelitinib, or gefitinib for 4 hours. **D)** Principal component analysis is shown for the top 500 most variable genes. **E - F)** Selected differentially expressed genes for 25 µM lapatinib (**E**) or 25 µM varlitinib (**F**) relative to 25 µM pelitinib and gefitinib conditions are shown. Genes are selected and annotated based on known functions. Variance-stabilized counts are z-scored on a per-gene basis across all samples.

To further characterize the effects of lapatinib and varlitinib, we cultured axenic Mtb in lapatinib- and varlitinib-containing media for 4 hours at near-MIC (5 µM) and above-MIC (25 µM) concentrations, normalized to 0.25% DMSO. Since we failed to detect effects on resazurin reduction or intracellular ATP for gefitinib or pelitinib, we used these conditions to control for transcriptional responses to the shared quinazoline/quinoline scaffold. We isolated and sequenced bulk RNA, then performed principal component analysis to visualize transcriptional similarity between Mtb samples exposed to each drug. Lapatinib-treated samples strongly separated from other samples and accounted for 85% of gene expression variability across the dataset (Figure 2D). This suggested that lapatinib and varlitinib have different transcriptional effects on axenic Mtb and potentially different mechanisms of action. We then performed differential expression analysis of lapatinib and varlitinib relative to reference drugs gefitinib and pelitinib and defined differentially expressed genes (DEGs) as the intersection of significantly differentially expressed genes (moderated |log_2_FC| > 1 in both conditions; p_adj_ < 0.05) relative to both reference drugs (Supplementary Figure 4).

At both doses, lapatinib differentially regulated genes associated with growth arrest and stress. Lapatinib treatment downregulated genes associated with ribosomal biogenesis (*rps* and *rpl* genes) and upregulated genes associated with oxidative stress (*furA, katG, cysD, cysN, cysO, cysM, mec, mshA, mshC, mshD*), DNA damage repair response (*recA, lexA, dnaE2, pafA, pafB, pafC, ruvA, ruvB, xthA, ung, lhr*),^26^ and the envelope damage-responsive operon *iniBAC* (Figure 2E). Lapatinib treatment also differentially regulated multiple groups of electron transport chain (ETC) genes, which may suggest metabolic rewiring upon acute lapatinib treatment. These transcriptional changes overlap with transcriptional effects of ETC-targeting respiration inhibitors, transcriptional inhibitors, fluoroquinolones, and cell wall synthesis inhibitors,^27–30^ and are suggestive of effects on bacterial physiology that might be linked to the antibiotic activity of lapatinib.

Compared to lapatinib, few differentially expressed transcripts were specifically associated with varlitinib treatment. Varlitinib induced genes from the lysine catabolic pathway (*lat),* pyrimidine biosynthesis pathway (*pyrB, pyrC, Rv1382*), efflux genes and their putative regulator (*mmpS4, mmpL4*, *Rv0452*), and *Rv0047c (napM)* (Figure 2F). Additional members of each pathway were significantly differentially regulated relative to one reference, but failed to reach the moderated log_2_ fold change threshold for both references. Induction of lysine catabolic genes has been associated with multiple non-replicative states,^31,32^ and lysine accumulation is thought to be toxic or act as an alarmone.^33,34^ Likewise, napM is induced by multiple stresses and inhibits DNA replication.^35^ This provides potential pathways to investigate and target during antibiotic development.

### Drugs without detected axenic antitubercular activity have dose-dependent effects on Mtb transcriptional state

Principal component analysis suggested that drugs without strong effects on Mtb growth phenotype, like high-dose gefitinib, might nonetheless alter bacterial transcriptional state. Although excluding lapatinib from principal component analysis allowed separation of varlitinib-treated samples from low-dose gefitinib- or pelitinib-treated samples, gefitinib and pelitinib conditions separated according to dose along the same principal components, and samples treated with high-dose gefitinib and high-dose varlitinib clustered together, suggestive of dose-dependent effects on transcriptional state (Figure 3A). Indeed, genes with high loadings on principal component 1 also were significantly differentially expressed in a dose-dependent manner upon gefitinib treatment (Figure 3B). Compared to varlitinib and lapatinib, which have strong effects on axenic Mtb at 25 µM, however, gefitinib and pelitinib led to dose-dependent differential expression of fewer genes (Figure 3C; Supplementary Table 1).

**Figure 3.**
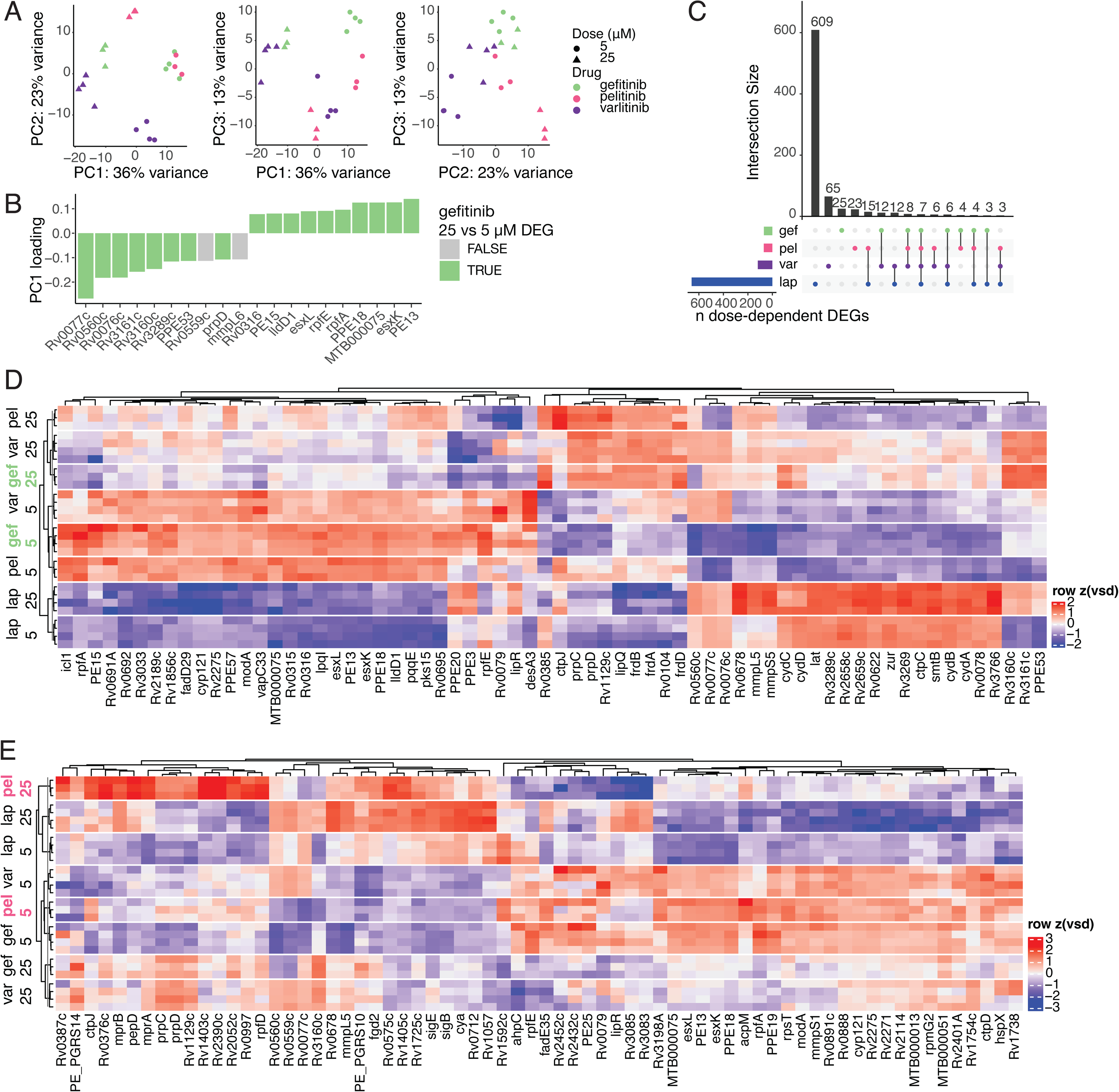
High-dose drug treatment has transcriptional effects, even for drugs without strong growth or metabolic effects. **A)** Principal component analysis of Mtb transcriptomes is shown for the top 500 most variable genes after exclusion of lapatinib-treated samples. The first three principal components, which collectively explain 72% variance, are plotted against one another. **B)** Top 20 loadings by magnitude for principal component 1. **C)** Between-condition overlap of genes with significant dose-dependent differential expression. **D - E**) Heatmap of all gefitinib-dose-dependent (**D**) and all pelitinib-dose-dependent (**E**) genes.

High-dose gefitinib treatment significantly upregulated gene sets suggestive of altered metabolism (Figure 3D). These metabolic gene sets included genes involved in propionate metabolism (*prpD, prpC, Rv1129c*), reduction of fumarate (*frdA, frdB, frdD)*, terminal oxidation by the alternative ETC cytochrome *bd* oxidase (*cydA, cydB, cydC, cydD)*, and the core lipid response (*Rv3160c, Rv3161c, PPE53*).^36^ A variety of *in vivo* and *in vitro* stresses slow aerobic respiration, leading to redox imbalances that are dissipated by transcriptionally regulated metabolic remodeling.^37–40^ This raises the possibility that high-dose gefitinib treatment directly affects bacterial metabolism. Pelitinib treatment also significantly differentially regulated Mtb gene expression in a dose-dependent manner. Pelitinib treatment significantly induced *marR* regulon members (*Rv1403c, Rv2390c, rpfD, Rv1405c*), the *mprAB-sigE-pepD* regulon, propionate metabolic genes (*prpC, prpD, Rv1129c*), and members of the *mymA/virS* operon (Figure 3E). While we classify pelitinib, poziotinib, and gefitinib as host-dependent and lapatinib and varlitinib as host-independent based on activity in broth and human macrophages, transcriptional profiling and moderately slowed growth kinetics suggests that direct effects on Mtb may contribute to the ostensibly host-dependent activities of pelitinib, poziotinib, and gefitinib.

### Host-dependent therapy is associated with differential expression of Mtb genes associated with environmental response

To investigate potential mechanisms of host-dependent growth restriction, we characterized Mtb transcriptional state under growth-restrictive, intramacrophage conditions. We reasoned that Mtb transcriptional state would be associated with bacterial restriction, as differences in intracellular Mtb state have been previously linked to restrictive or permissive host cell subsets and phagosomal environments.^41,42^

Pelitinib and gefitinib have different scaffolds and off-target effects. Since pelitinib is a 4-aminoquinoline EGFR inhibitor with off-target and weaker activity against Src kinase,^43^ we considered that differences in scaffold or Src inhibition could lead to different restriction phenotypes by pelitinib and gefitinib. As a growth-restrictive reference condition, we used the drug saracatinib, which inhibits Src kinase, but has a 4-aminoquinazoline scaffold in common with gefitinib (Figure 4A). We infected primary human macrophages derived from 6 donors with H37Rv Mtb, then treated infections with 10 µM pelitinib, gefitinib, saracatinib, or vehicle for 24 hours before RNA isolation, polyA-independent library preparation, and hybrid selection-based enrichment of Mtb-derived library fragments (Figure 4B). In parallel, we tracked Mtb luminescence over time for each treatment condition. At 24 hours post infection, Mtb inside drug-treated macrophages had lower luminescence (Figure 4C), indicative of intracellular restriction. Consistent with this phenotype, we detected 50, 62, and 208 differentially expressed Mtb genes (|moderated log_2_FC| > 1; p_adj_ < 0.05) relative to vehicle for pelitinib, gefitinib, and saracatinib conditions respectively (Figure 4D).

**Figure 4.**
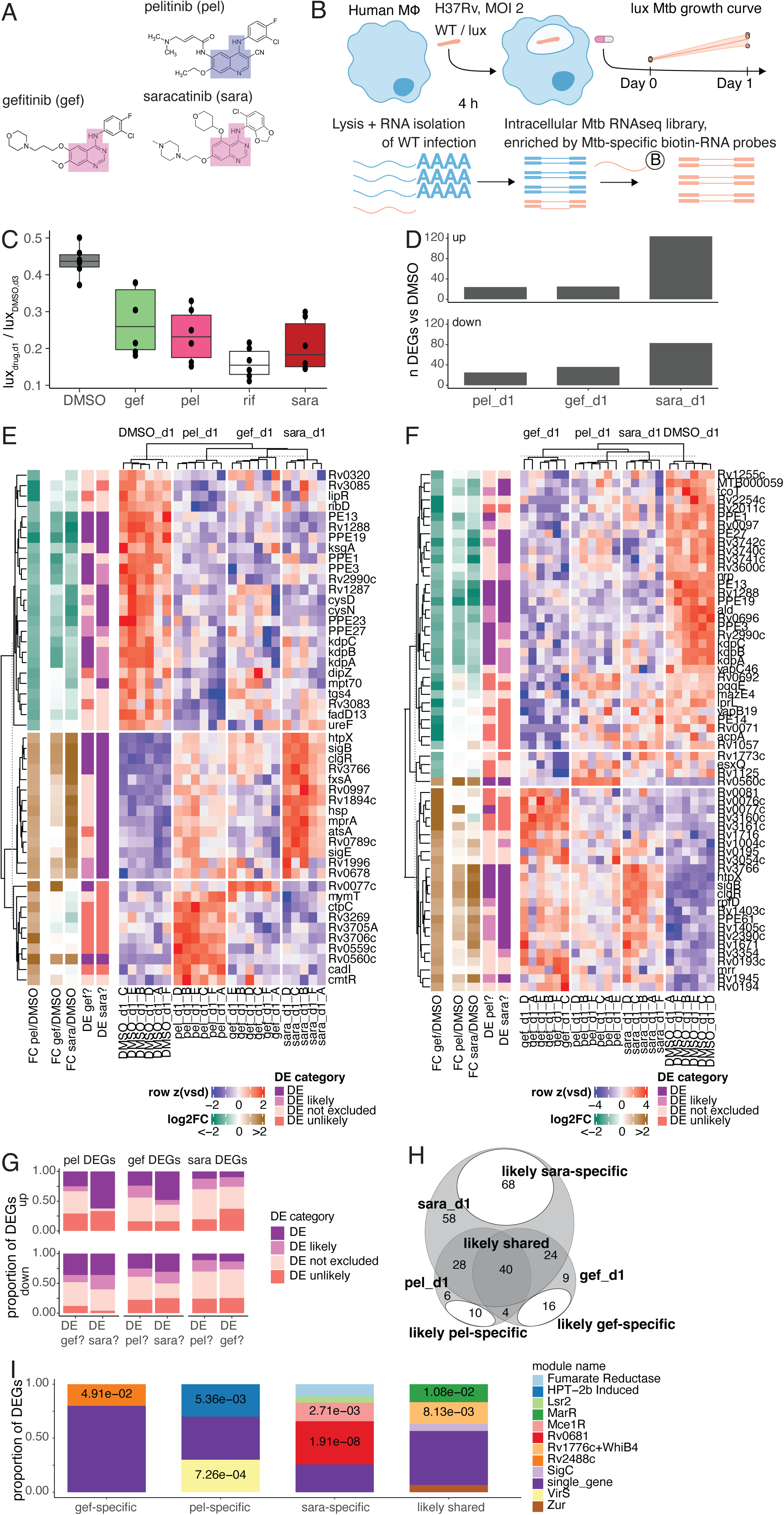
Host-dependent therapies drive heterogeneous, partially overlapping combinations of environment-associated Mtb transcriptional responses. **A)** Chemical structure of pelitinib, gefitinib, and saracatinib. The 4-aminoquinoline and 4-aminoquinazoline scaffolds are highlighted in blue and pink, respectively. **B)** Experimental schematic of parallel infections for intracellular luminescence measurement and RNA isolation. **C)** Day 1 intracellular Mtb luminescence in macrophages treated with 10 µM drug, rifampicin, or DMSO, normalized to DMSO luminescence at day 3. Points represent the mean of triplicate infections for n=6 donors. **D)** Number of differentially expressed Mtb genes (|moderated log_2_ FC| > 1, padj < 0.05) at 24 hours post-infection for each drug relative to DMSO. **E - F)** Differentially expressed Mtb genes in pelitinib-treated macrophages (**E**) and gefitinib-treated macrophages (**F**) relative to DMSO-treated macrophages at 24 hours, z-scored variance-stabilized counts for n=5-6 donors. Genes are annotated with moderated log_2_ fold change estimates for each drug relative to DMSO and statistically categorized according to differential expression status in other drug conditions relative to DMSO. **G)** Summary categorization of each drug’s DEGs by differential expression status in the other two treatment conditions. **H)** Overlap in likely DEGs relative to DMSO across pelitinib, gefitinib, and saracatinib conditions. Likely drug-specific genes within non-overlapping regions of the Venn diagram are highlighted. **I)** Enrichment of iModulonDB-defined Mtb gene modules^50^ in shared and drug-specific gene sets. Benjamini-Hochberg–corrected p values for significant hypergeometric tests using the background set of all DEGs (n=320) are reported.

For each condition, changes to transcript expression included upregulation of two-component systems, sigma factors, transcription factors, and environment-responsive genes (Figure 4E-F; Supplementary Figure 5; Supplementary Figure 6). Pelitinib-treated samples were characterized by differential upregulation of genes induced by divalent metal cations (*mymT, cadI, cmtR, Rv1996, ctpC, Rv3269*). We also observed downregulation of the low-potassium-responsive Kdp potassium transport system (*kdpA, kdpB, kdpC*) and a majority of the *mymA/virS* operon (*Rv3083 (mymA), lipR (Rv3084), Rv3085, tgs4 (Rv3088), fadD13 (Rv3089)*). Gefitinib-treated samples were instead characterized by upregulation of acid-responsive *marR* regulon members (*Rv0194, Rv0195, PPE61, Rv1403c, Rv1405c, Rv2390c*)^44^ and downregulation of low-potassium-responsive genes (*kdpA, kdpB, kdpC, PE27*).^45^ Saracatinib-treated samples were characterized by downregulation of mce1 operon members (*yrbE1A, yrbE1B, mce1A, mce1B, mce1C, mce1D, lprK, mce1F*) and upregulation of genes related to kstR-regulated cholesterol catabolism (*kstR, cyp142, hsaD, fadE26, fadE34, kshA, Rv3502, Rv3530c, Rv3531c*), as well as marR regulon genes (*Rv0575c, Rv1403c, Rv1405c, Rv2390c, PPE61*).

Mtb DEGs for each drug condition displayed two major kinetic behaviors at 24 hours compared to a post-phagocytosis, pre-treatment reference (Supplementary Figure 7). Besides being differentially expressed between drug and vehicle at 24 hours, one subset of DEGs was induced or repressed post-phagocytosis in both conditions, suggestive of these genes’ role in early transcriptional adaptation more generally. Another subset was exclusively induced or repressed at the 24 hour timepoint in drug-treated macrophages but had unchanged expression post-phagocytosis in vehicle-treated macrophages, suggesting that host-dependent therapy induces environment-associated Mtb responses in addition to tuning expression magnitudes of genes involved in early transcriptional adaptation. Together, we show that intracellular Mtb respond to different stimuli in drug-treated macrophages relative to vehicle-treated macrophages, with each host-dependent drug driving distinct Mtb environmental responses.

### Environment-responsive Mtb genes can be statistically categorized as shared among growth-restrictive contexts or unique to gefitinib, pelitinib, or saracatinib treatment

Comparison of each drug’s DEGs suggested that growth restriction by each drug is associated with both a general, shared Mtb stress response and a drug-specific environmental response (Supplementary Table 2). However, strictly comparing DEGs above a fold-change cutoff likely underestimates shared genes and overestimates drug-specific genes, as cutoff-based approaches exclude genes with moderate but robust changes in expression and do not distinguish true low-change genes from noisy genes with low-confidence fold change estimates. To compare Mtb DEGs for two treatment conditions relative to vehicle, we used a Wald test to determine whether one test condition’s DEGs had negligible unmoderated fold changes in the second test condition and were therefore unlikely to be differentially expressed (Supplementary Figure 8A). Remaining genes were categorized as likely differentially expressed if they had a moderated fold change above the Wald test threshold, or as unspecified. Since threshold-based statistical tests are more conservative than post-hoc thresholds (Supplementary Figure 8B),^46^ we empirically tested a range of log_2_ fold change thresholds and used 0.7 as a tradeoff between stringency and categorization of genes across a range of expression levels (Supplementary Figure 8C-D).

Using this statistical framework, we categorized pelitinib, gefitinib, and saracatinib DEGs as differentially expressed, likely differentially expressed, unlikely differentially expressed, or unspecified in other conditions. Statistical categorizations were validated by clustering DEG expression patterns across all drug conditions (Figure 4E-G; Supplementary Figure 6). Pelitinib-downregulated DEGs formed a cluster that included genes with shared differential expression, as well as genes that could not be confidently specified as pelitinib-specific or shared, while pelitinib-upregulated DEGs formed two distinct clusters according to pelitinib-specificity (Figure 4E). One cluster of pelitinib-upregulated DEGs consisted of sigE-regulated genes^47,48^ and had entirely shared differential expression between pelitinib and saracatinib treatments, with a smaller subset of these genes shared across all three treatments. The second cluster of upregulated genes was largely pelitinib-specific and corresponded to divalent metal cation-responsive genes and *tcrXY* target genes *Rv3705A* and *Rv3706c*.^49^ Within gefitinib and saracatinib DEGs, groups of genes associated with a specific environmental response also generally clustered together, and genes within a group displayed similar patterns of drug-specificity (Figure 4F; Supplementary Figure 6). Together, this suggests that Mtb DEGs associated with each growth-restrictive treatment belong to coherent gene sets, which are defined both by their drug-specificity and by genes associated with distinct environmental responses and regulators.

To compare transcriptional states across conditions, we used the set of likely differentially expressed genes, which allowed us to define shared genes more permissively without entirely relaxing differential expression criteria (Figure 4H; Supplementary Figure 9A-D). The intersection set comprised 40 genes with likely shared differential expression across all growth-restrictive conditions. We also defined 10, 16, and 48 drug-specific genes for pelitinib, gefitinib, and saracatinib conditions, respectively.

### Mtb pathway enrichment supports a model where host-dependent therapies induce different combinations of environment-responsive pathways

We hypothesized that shared Mtb genes could reveal shared mechanisms of or transcriptional adaptations to growth restriction, while drug-specific genes could reveal biological differences in the Mtb stress response across conditions. To relate shared and drug-specific genes to known regulatory pathways, we tested hypergeometric-based enrichment of independent gene modules derived from machine-learning–based decomposition of an Mtb gene expression compendium.^50^ Mtb genes with shared differential expression were significantly enriched for modules annotated as MarR-regulated and Rv1776c+WhiB4-regulated (Figure 4I; Supplementary Figure 9A; Supplementary Figure 10A-B), with the Rv1776c+WhiB4 module partially recapitulating genes in the literature-defined sigE regulon.^47^ In addition to being enriched among the list of shared Mtb genes, MarR- and Rv1776c+WhiB4/sigE-regulated genes were also significantly and separately enriched in each condition relative to vehicle (Supplementary Figure 10C-H). These shared Mtb transcriptional responses to pelitinib-treated, gefitinib-treated, and saracatinib-treated macrophages may reflect core Mtb adaptation strategies during growth restriction or core features of a growth-restrictive host environment.

Pelitinib-specific genes were significantly enriched for two modules annotated as HPT-2b–induced and VirS-regulated (Supplementary Figure 9B). HPT-2b is an antitubercular agent that drives intracellular copper accumulation and copper toxicity-associated transcriptional responses.^51^ Enrichment of the HPT-2b–induced module may suggest Mtb in pelitinib-treated macrophages experience a unique environmental stress that has features in common with metal toxicity. Gefitinib-specific genes largely did not map to particular modules, aside from the enriched Rv2488 module, which consists of a “core lipid response”^36^ that is also induced by several antibiotics (Supplementary Figure 9C).^52–55^ Since few environment-associated Mtb gene modules were uniquely induced in gefitinib-treated macrophages, this suggests that Mtb stresses associated with gefitinib treatment are largely a subset of environmental stresses associated with pelitinib or saracatinib treatments. However, it is also possible that gefitinib-specific genes could have expression kinetics that lead to most members of a module falling outside of differential expression thresholds at the 24 hour timepoint.

For saracatinib-specific Mtb genes, Mce1R-annotated and Rv0681-annotated modules were significantly enriched (Supplementary Figure 9D; Supplementary Figure 10I). Saracatinib-specific Mtb genes in the Mce1R-annotated module were downregulated relative to vehicle, while those in the Rv0681-annotated module were upregulated. The Mce1R-annotated module contains most members of the *mce1* operon, which encodes a complex required for fatty acid import.^56^ Since the Rv0681-annotated module contains genes encoding almost all enzymes required for cholesterol oxidation, side-chain degradation, and ring opening, as well as their regulator *kstR*,^57,58^ this module partially recapitulates the *kstR* regulon and likely reflects a shift toward cholesterol catabolism. Gene set enrichment analysis using the experimentally defined *kstR* regulon further validated a saracatinib-specific cholesterol catabolic signature.^58^ (Supplementary Figure 10J-L). Concomitant downregulation of *mce1* genes and upregulation of *kstR*-regulated genes could be consistent with an overall shift in Mtb lipid metabolism, either as a metabolic adaptation to environmental stress or as a reflection of the host carbon source environment. Together, we find that Mtb inside pelitinib-treated, gefitinib-treated, and saracatinib-treated macrophages are in distinct transcriptional states characterized by partially overlapping combinations of environment-associated genes. In particular, host-dependent growth restriction by pelitinib, gefitinib, and saracatinib involves distinct Mtb stress responses, suggesting distinct and independently inducible mechanisms of intracellular growth restriction by each drug.

### Host-dependent therapy alters the transcriptional state of infected host cells

Despite a shared *sigE/Rv1776c+whiB4* and *marR* Mtb response across drug treatments, timepoint-matched host transcriptomes did not clearly indicate transcriptional activation of a shared, causative antimicrobial pathway (Supplementary Figure 11). Innate immunity-related transcripts (*TLR2, CXCL8, IL1B, IL1RN, IL18, CCL3, CCL4*) were downregulated, consistent with altered cytokine signaling described in gefitinib-treated, Mtb-infected murine macrophages.^13^ Cholesterol efflux-related genes (*ANXA6, EEPD1, OLR1, ABCA8*),^59–62^ which might relate to phagosomal acidification,^63^ also had shared differential expression. Pelitinib-treated macrophages were also characterized by antimicrobial-associated transcripts not induced by gefitinib, such as *GPR183*,^64^ *HAMP*,^65^ and zinc-binding metallothionein *MT1G.*^66^

### Differential gene expression in pelitinib- and gefitinib-treated macrophages consists of host environment-dependent and host-independent components

Since EGFR kinase inhibitors are sufficiently membrane-permeable to target cytoplasmic kinase domains of EGFR, they may also access the phagosome to act directly on Mtb. To identify transcriptional effects that are potentially due to direct drug-Mtb interaction, rather than host environment-dependent effects, we sequenced RNA from Mtb cultures grown in 10 µM pelitinib-, gefitinib-, or vehicle-containing 7H9 broth for 24 hours.

Using the same fold change and significance thresholds, we identified 18 and 5 axenic differentially expressed genes for pelitinib and gefitinib treatments, respectively, of which a subset were also differentially expressed intramacrophage (Figure 5A; Supplementary Figure 12A). There was not a strong relationship between log_2_ fold change in axenic and intramacrophage experiments, and we observed genes with significant axenic differential expression but low intramacrophage log_2_ fold change estimates and *vice versa* (Figure 5B).

**Figure 5.**
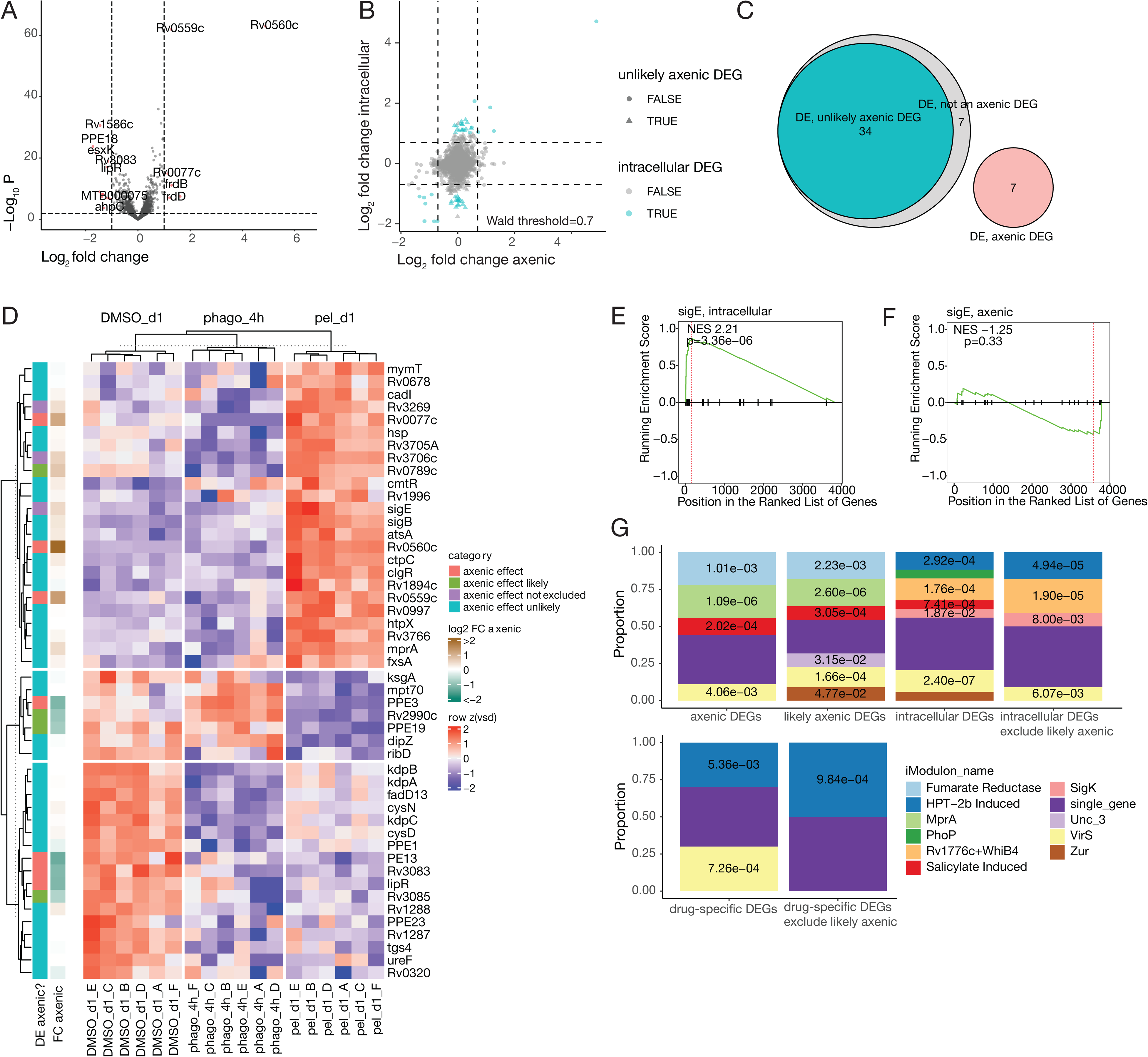
Effects of pelitinib therapy on Mtb gene expression include host environment-dependent and axenic effects. **A)** RNA sequencing of axenic Mtb cultures exposed to 10 µM pelitinib or vehicle in 7H9 for 24 hours, n=4 biological replicates. Log_10_ Benjamini-Hochberg–adjusted p values are plotted against moderated log_2_ fold change estimates for each gene, and genes with significant differential expression are colored in red. **B)** For each gene, moderated log_2_ fold changes for intracellular and axenic conditions are plotted against each other. Genes are colored according to their intracellular differential expression and plotted as triangles if axenic differential expression is unlikely (Wald test |FC| < 0.7, padj < 0.05). **C)** Overlap in categorization of intracellular DEGs as unlikely to be axenically differentially expressed by Wald test (unlikely axenic effect), not determined to be axenically differentially expressed by post-hoc threshold (not an axenic DE), and axenically differentially expressed (axenic DE). **D)** Heatmap of intracellular differential expression in pelitinib relative to DMSO at 24 hours, annotated according to whether genes are statistically categorized as exhibiting axenic differential expression. **E - F)** Gene set enrichment analysis of literature-defined sigE regulon genes^47^ in intracellular (**E**) and axenic (**F**) condition using the list of all genes ranked by moderated log_2_ fold change. **G)** Hypergeometric enrichment of iModulonDB-defined Mtb gene modules^50^ in pelitinib and pelitinib-specific DEGs, excluding genes with axenic differential expression. For pelitinib DEGs, the background set is defined as all iModulon-mappable genes, while for pelitinib-specific DEGs, the background set is defined as all DEGs.

Using a Wald test against a log_2_ fold change threshold of 0.7, we could define 34 intramacrophage pelitinib DE genes as unlikely to be axenically differentially expressed, 7 as unspecified, and 7 intramacrophage DE genes as axenically differentially expressed (Figure 5C-D). For gefitinib DE genes, we similarly did not observe a relationship between axenic and intramacrophage log_2_ fold change estimates. Given that sigE is differentially expressed in pelitinib-treated macrophages and we failed to rule out axenic differential expression, we tested whether we could detect modest upregulation of the sigE regulon by gene set enrichment analysis (Figure 5E-F). While pelitinib treatment significantly enriches for sigE regulon genes in intramacrophage Mtb, the sigE regulon is not significantly enriched in axenic Mtb at the same timepoint. This suggests that our approach is conservative in its identification of axenic, host-independent effects.

Because axenic drug stimulation was sufficient for induction of a subset of DEGs in drug-treated macrophages, we hypothesized that some differences in intramacrophage Mtb environmental response between drug treatments could be due to differences in phagosomal Mtb-drug interactions. While enrichment of the *mymA/virS* operon was stronger in pelitinib-treated macrophages than pelitinib-treated Mtb cultures, we could partially attribute intramacrophage enrichment of the virS module to axenic expression of *Rv3083, lipR,* and *Rv3085* (Figure 5D). The gefitinib-specific Rv2488 module could be entirely attributed to axenic expression of *Rv3160c* and *Rv3161c* (Supplementary Figure 12B-C). This suggests that a subset of the drug-specific transcriptional response could be due to presence of the drug in the intraphagosomal environment, leading to direct transcriptional effects. Although it is possible that particular host environment-dependent effects require drug-Mtb effects, we provide a framework for categorizing Mtb-drug-host interactions and show pelitinib and gefitinib have host environment-dependent effects beyond their effects in axenic culture.

Together, we show that host-dependent therapies are associated with several possible intracellular Mtb states, which can be decomposed into environment-associated modules and contributions from host-independent, drug-Mtb interactions and from host environment-dependent effects.

## Discussion

Tuberculosis treatment involves lengthy antibiotic regimens, risking resistance development. Enhancing host antimicrobial processes could be a mode of host-directed therapy, complementary to antibiotic treatment. Despite the specific goal of enhancing antimicrobial processes, HDT discovery involves screening for intracellular growth restriction in general, excluding antibiotic hits, then identifying potential host-directed effects. Previous work has failed to isolate HDT-resistant mutants,^13^ although enhancing antimicrobial processes may itself drive antibiotic and stress tolerance.^67^ Measuring bacterial transcriptional state can resolve features of growth restriction in greater detail and, in turn, inform screening strategies and adjunctive use of HDTs during antibiotic therapy.

Using EGFR kinase inhibitors as a prototype, we measured Mtb restriction, Mtb transcriptional state, and host transcriptional state. Few tested EGFR inhibitors strongly restricted intracellular Mtb, and we failed to detect EGFR surface expression in our infection model. Growth-restrictive drugs fell along a spectrum of direct anti-Mtb effects, from host-dependent drugs with effects on axenic transcription but not phenotype to host-independent antibiotics with strong effects on axenic phenotype. This argues against a purely host-directed, EGFR-centric model of growth restriction. By comparing host-dependent drugs, we identify environmental stress responses associated with growth restriction, including WhiB4+Rv1776c and MarR gene modules (shared), HTP-2b–induced, metal toxicity-associated genes (pelitinib), and cholesterol-associated Mce1R and Rv0681 gene modules (saracatinib). These Mtb stress responses could act as transcriptional markers for specific antimicrobial states during screen design or mechanistic follow-up.

Many EGFR inhibitors are quinazolines, a chemical class that contains antimycobacterials.^21^ However, antibiotics, host-synergizing therapies, and host-directed therapies can be difficult to distinguish. Bedaquiline, an antimycobacterial ATP synthase inhibitor, also enhances phagosomal acidification.^68^ Antibiotics can also depend on the host environment without directly altering host state. Host stress drives Mtb adaptation, which can introduce antibiotic-exploitable vulnerabilities^52,69–72^ or cross-protect against antibiotics.^73,74^ We categorized varlitinib and lapatinib as host-independent, but varlitinib and lapatinib could also alter host state directly or synergize with the host environment.

We categorized pelitinib, gefitinib, and poziotinib as host-dependent as they lacked strong axenic growth or metabolic defects, but direct drug-Mtb interaction could contribute to host-dependent restriction. *Rv3160c-Rv3161c*, *Rv0077c-Rv0078*, and *Rv0559c-Rv0560c* were induced both axenically and intracellularly by gefitinib and pelitinib. These genes are antibiotic-induced,^52,55^ alter susceptibility to some but not all antibiotic and host-imposed stresses,^52,53,55,75,76^ and are mutated in antibiotic-resistant mutants, including quinazoline-resistant mutants.^77,78^ Mtb also axenically downregulates *mymA/virS* operon members *Rv3083, lipR,* and *Rv3085* in response to pelitinib, which may impact intracellular fitness, as *mymA/virS* is essential *in vivo* and in acidic conditions.^79,80^ Measuring axenic growth across diverse media conditions could clarify how pelitinib and gefitinib act.

Characterizing putative HDTs typically involves a top-down approach, identifying host-directed effects associated with intracellular growth restriction. We take a bottom-up, pathogen-focused approach, associating intracellular growth restriction with effects on bacterial state. Bacterial transcriptional state provides a relatively direct, holistic metric of how drug treatment affects intracellular bacteria. However, modeling how host antimicrobial state drives the bacterial states identified by our approach poses additional challenges. First, the phagosome is formed *de novo* and progressively recruits antimicrobial effectors from multiple subcellular compartments. Capturing host antimicrobial capacity involves measuring phagosomal ionic composition, pH, redox state, enzyme and antimicrobial peptide activity, and nutrient composition. Although transcriptomic methods scale well, the degree of transcriptional regulation involved in phagosome biochemistry is not well understood. Second, in the host-directed hypothesis, acquisition of host antimicrobial function causes bacteria to adopt a stressed state. Modeling causal interactions between host and pathogen states requires assumptions about which host and pathogen timepoints to correlate and how to incorporate feedback from pathogen to host. Large-scale, high-dimensional data collection will be critical to developing host-pathogen interaction models with biologically valid assumptions.

Mtb transcriptional state is a proposed readout of the phagosomal environment.^41^ Several Mtb transcriptional regulators sense stress and are positively regulated by stress exposure, allowing their differential expression to act as an environmental proxy. Axenic studies have defined modules of stress-responsive genes, but it is difficult to infer the exact phagosomal stress that caused upregulation of a particular gene, given regulatory cross-talk, anticipatory responses, and shared adaptation strategies to unique stresses.^81–84^ Despite this, we observed distinct, environment-responsive modules upregulated in gefitinib-, pelitinib-, and saracatinib-treated macrophages. In vehicle-treated macrophages, Mtb upregulated genes encoding the low potassium-inducible Kdp potassium transporter system. In all drug conditions, Mtb upregulated *marR* regulon genes, which are pH- and chloride-sensitive,^44,85^ and downregulated Kdp subunit-encoding genes. This could reflect responses to different phagosomal environments, since the phagosome acidifies and acquires chloride and potassium during maturation.^45,85^ In pelitinib- or saracatinib-treated macrophages, additional stress modules were differentially regulated, such as genes typically induced by envelope stress, divalent metal cations, and changes to cholesterol availability. This could suggest multiple, independently inducible routes to Mtb restriction in human macrophages. This multi-stress model of HDT-inducible restriction in human macrophages differs from the single-stress model of interferon gamma-inducible control in mice, which is almost entirely dependent on inducible nitric oxide synthase.^41^ Detailed characterization of phagosome states during treatment can help clarify how transcriptional signatures relate to host-imposed stresses.

Although transcriptional state reflects an intermediate layer between DNA and protein expression, transcript-protein concordance is gene- and context-dependent. During nitrosative stress, changes to transcript and protein expression are uncoupled.^86^ Acute transcriptional changes take hours to be reflected at the protein level, and acute proteomic changes involve targeted protein degradation rather than translation from *de novo* transcripts. Codon-biased translation during stress can also decouple transcript and protein levels.^87^ Differentially abundant transcripts may not reflect machinery required to detoxify acute stress. Instead, differentially abundant transcripts define Mtb states within a general, growth-restricted phenotype, provide a short half-life measurement of stress responses, and may reflect longer-term adaptations required to survive stress or restore homeostasis. Future work could link transcript-level changes to intracellular fitness and precise mechanisms of restriction.

When intramacrophage drug screens identify growth-restrictive drugs, these drugs should be designated as “host-directed” with caution and validation should equally account for alternative mechanisms of action.

Since EGFR kinase inhibitors emerge from screens for intracellular Mtb growth restriction,^3–5^ one hypothesis is that EGFR is a host factor regulating infection outcome. Small molecule screens can identify infection-regulating host kinases while accounting for polypharmacology by regressing target profiles against an outcome of interest.^88–90^ However, several of the EGFR inhibitors we screened had high reported selectivity (lapatinib, varlitinib, afatinib, canertinib). While inhibitor-kinase target profiles cover the human kinome,^91–93^ similar profiles do not exist for the Mtb kinome. When eukaryotic kinase inhibitors are antibiotic *via* inhibition of essential Mtb eukaryotic-like serine/threonine protein kinases,^94^ they are excludable using axenic counter-screens. However, off-target inhibition of non-essential Mtb kinases may not inhibit axenic growth but may alter intracellular fitness. For example, the Mtb osmosensing kinase PknD is inhibited by SP6000125, which was originally developed as a JNK inhibitor.^95^ Confirming expression of targetable host kinases and inhibitor selectivity for pathogen and host kinases can aid interpretation of screen hits.

Defining a repurposed drug as “host-directed” is a mechanistic hypothesis about where effects on intracellular growth restriction arise. Because intracellular growth restriction is the result of bidirectional, host-pathogen interactions, the host-directed hypothesis can be difficult to test, and alternative hypotheses can be difficult to exclude.

First, host-directed effects must be identified, perturbed, and defined as necessary and sufficient for intracellular growth restriction. A common perturbation strategy involves using small molecules that inhibit canonical phagosome functions, thereby identifying host processes required for growth restriction. However, due to polypharmacology of small molecule inhibitors and the coupled nature of phagosome stresses,^96–98^ it is difficult to perturb phagosome state in a targeted manner or identify the small molecule-disrupted process responsible for growth restriction without re-measuring phagosome state. Panels of beads functionalized with different probes of phagosomal biochemistry could be deployed during characterization of screen hits or validation of small molecule perturbations. Alternatively, parallel screening with probe-functionalized beads and bacteria would ensure screen hits have host-directed effects on baseline antimicrobial capacity and alter bacterial restriction during infection.

Second, bacteria-directed effects must not exist or be unrelated to growth restriction. Direct antibiotic effects are tested by measuring a drug’s effect on axenic phenotype, e.g. growth, survival, or metabolic activity. If a host-directed therapy restricts intracellular growth at a specific concentration but does not affect axenic phenotype at the same concentration, it is sometimes assumed to be primarily host-directed. However, the minimum inhibitory concentration of *bona fide* antibiotics is environment-dependent,^52,69–72^ and the intracellular concentration of a drug can be orders of magnitude higher than its extracellular concentration.^99^ Confident exclusion of antibiotic effects therefore relies on choice of screening environment and concentration. Since intracellular Mtb are thought to reside in an acidic, cholesterol-rich phagosomal environment, antibiotics with pH-dependent or cholesterol-dependent activity may initially appear host-directed. If bacteria-directed effects are undesired, stringent testing for antibiotic effects at doses higher than the treatment dose is required.

Finally, excluding bacteria-directed effects involves excluding effects on the host *via* effects on the bacterium; for example, drug treatment could allow the host cell to access a new antimicrobial state by preventing Mtb effector secretion.^100^ High-throughput sequencing of Mtb transcriptomes^101^ or fluorescent transcriptional reporters can discover bacteria-directed effects not captured by axenic phenotype.

While these distinctions do not change the therapeutic potential of drug repurposing, precise classification of repurposed drugs can guide their optimization as therapeutics and inform how we relate small molecule perturbations to the biology of host-pathogen interactions.

## Materials and methods

### Bacterial culture

*Mycobacterium tuberculosis* strain H37Rv was grown in Middlebrook 7H9 supplemented with 10% OADC (Fisher, cat #BD212351), 0.2% glycerol, and 0.05% Tween-80. Strains containing plasmids were grown in the presence of appropriate antibiotic: Hygromycin B (50ug/mL; Sigma cat #10843555001) and zeocin (20ug/mL; Thermofisher cat #R25001).

### Macrophage isolation and culture

Deidentified buffy coats were collected from healthy human donors at Massachusetts General Hospital. CD14+ monocytes were isolated from buffy coats by density-based centrifugation in Ficoll Paque Plus (Millipore Sigma, cat #GE17-1440-03), followed by CD14+ positive selection (Stemcell, cat #17858). Monocytes were differentiated 6 days in RPMI (Thermofisher, cat# 11835030) supplemented with 10% FBS (Thermofisher, cat #10082147), 1% HEPES (Corning, cat #25-060-CI), 2 mM L-glutamine (Sigma, cat #G7513), and 25 ng/mL M-CSF (Biolegend, cat #574804). After differentiation, macrophages were maintained in supplemented RPMI without M-CSF.

### Macrophage infections

Mid-log phase bacteria were pelleted at 3900 rpm, washed once in DPBS, resuspended in cell culture media, then passed through a 5 um filter to generate a single cell suspension. For lux experiments, bacteria expressing an integrating luxABCDE cassette were used. For RNA sequencing experiments, rather than filtering bacteria, bacteria were spun at 500 rpm to settle larger clumps. Macrophages were infected at MOI 2 for 4 hours, washed three times in DPBS, and then cultured in the appropriate media.

For experiments assessing drug effect on intracellular bacteria, 50,000 macrophages were plated in tissue culture-treated, white-bottom, 96-well plates (Corning, cat #353377) 24 hours prior to infection. Drugs were added after the infection period at a standard concentration of 10 µM unless otherwise noted, and intracellular luminescence was measured daily using a Tecan Spark 10M plate reader.

### Surface flow cytometry for EGFR and ErbB family receptors

Cells were detached from low-attachment plates using 2 mM EDTA in DPBS. Detached cells were pelleted at 500xg, blocked in TruStain FcX (Biolegend, cat #422302) 10 minutes at room temperature, before staining with PE-conjugated anti-ErbB receptor antibodies or isotype (Biolegend: anti-EGFR cat #352903, anti-ErbB2 cat #324406, anti-ErbB3 cat #324705, anti-ErbB4 cat #324706, isotype IgG1 K cat #981804, isotype IgG2a K cat #981910) at 1:50 for 30 minutes. After staining, cells were fixed and analyzed on a BD LSRFortessa cytometer.

### Protein isolation and western blotting for phospho- and total EGFR

For protein isolation, 500,000 A549 cells or human monocyte-derived cells were plated in tissue culture-treated, 12-well plates. Cells were stimulated with recombinant human epidermal growth factor (Biolegend, cat #585506) at 100 ng/ml or physiological water as a carrier control for 15 minutes before lysis in 1X RIPA buffer containing phosphatase inhibitor (Thermo Scientific), HALT protease inhibitor (Thermo Scientific), and 1% SDS. Lysates were treated with benzonase at 1:1000 (Sigma, cat #70664-3) for 30 minutes at room temperature, then clarified for 10 minutes, 12,000g. Protein concentration in clarified lysates was quantified by Pierce 660 nm assay (Thermo Scientific).

Each lysate sample was boiled in 1X Laemmli buffer for 5 minutes at 95C. For each sample, 17 ug lysate was loaded onto a 4-12% Bis-Tris gel, then transferred onto a nitrocellulose membrane using the iBlot 2 dry blotting system (Invitrogen). Membranes were blocked at room temperature for 1 hour in Intercept TBS Blocking Buffer (LICORbio, cat #927-60001). Blocked membranes were then stained overnight with primary antibodies (Cell Signaling Technology: rabbit EGFR antibody cat #4267, rabbit phospho-EGFR antibody cat #3777; Biolegend: rat beta-actin antibody cat #664801) diluted 1:1,000 in Intercept TBS Antibody Diluent (LICORbio, cat #927-65001) at 4C. Primary-stained membranes were washed thrice in TBS-T, incubated with fluorescent secondary antibodies (LICORbio: IRDye 680RD goat anti-rabbit IgG cat #926-68076, IRDye 800CW goat anti-rat IgG cat #926-32219) at a 1:10,000 dilution for 1 hour at room temperature. Secondary-stained membranes were washed thrice in TBS-T before imaging on a Licor Odyssey CLx imager.

### Resazurin conversion assay

Mid-log phase bacteria were diluted to OD 0.003 and grown in drug-containing 7H9 for 4 days before addition of Alamar blue (BioRad) to a final concentration of 10% v/v. Vehicle concentration in all conditions was 0.5% DMSO. Bacteria were incubated an additional two days before absorbance measurements using a Tecan Spark 10M plate reader.

### Intracellular ATP measurement

Mid-log phase bacteria were diluted to OD 0.1 and exposed to 5 or 25 µM drug for 24 hours. All concentrations were normalized to constant vehicle concentration of 0.25% DMSO (v/v). The optical density of drug-treated bacteria was measured, then 1 mL aliquots were heat-killed for 30 minutes at 95C. 25 µL heat-killed bacteria were incubated with an equal volume of BacTiterGlo (Promega) in flat white 96 well plates before luminescence measurements using a Tecan Spark 10M plate reader. ATP content per bacterial cell was calculated relative to DMSO by normalizing relative luminescence to input OD, then to normalized luminescence for DMSO.

### Intracellular and axenic bacterial RNA sequencing and quantification

Bacterial RNA was harvested from infected macrophages using a previously described differential lysis protocol.^42^ Infected macrophages were lysed in Trizol (Thermo Fisher Scientific) and centrifuged at max speed 4C for 20 minutes to pellet intact bacteria. The supernatant, representing the host RNA fraction, was collected and filtered twice through a 0.2 um PVDF filter for inactivation. To extract the bacterial RNA fraction, additional fresh Trizol and Lysing Matrix B (MP Bio) were added to the bacterial pellet. Bacteria were bead-beat on a benchtop homogenizer (BeadBug; Benchmark Scientific) at 400 rpm for 45 seconds, rested on ice for 5 minutes, followed by an additional 30 seconds of bead-beating. After bead-beating, the host RNA and bacterial RNA fractions were combined, organically extracted using 200 ul chloroform per 1 mL Trizol, then purified and DNAse-treated on-column (RNA Clean and Concentrate; Zymo).

200 ng isolated RNA was used as input for strand-specific library preparation using the Agilent SureSelect XT HS2 RNA Kit and their provided protocols. For intracellular bacterial RNA sequencing, 200 - 1000 ng unpooled libraries were hybrid selected for fragments originating from bacterial transcripts using a custom set of biotinylated probes complementary to the H37Rv transcriptome, excepting ribosomal and transfer RNAs. To account for the lower abundance of on-target bacterial transcripts relative to off-target host transcripts, probes were used at a 1:5 dilution. For axenic libraries, no hybrid selection was performed. All libraries were sequenced on NextSeq 550 (Illumina) using 150 - 300 bp read length.

Sequenced reads were quality-checked, aligned, and quantified using nf-core’s dualrnaseq pipeline v1.0.0, which we modified to include UMI deduplication by umi-tools v1.0.1. Prior to mapping, UMIs from each mate of the read pair were combined and extracted into each mate’s header using umi-tools extract. UMI-extracted paired-end reads were used as input for the mapping stage of the pipeline, where reads were mapped with STAR to a composite pathogen-host genome generated by the dualrnaseq pipeline from human genome version GRCh38 p13 and Mtb genome version NC_000963. The mapped file was indexed using samtools, deduplicated with umi-tools dedup, then quantified using HTSeq. For hybrid selected libraries only, libraries were sequenced and aligned in single-end mode, which precluded UMI-deduplication.

For paired host RNA library prep from infected macrophages, 20 ng isolated RNA was used as input for poly-A dependent library preparation. Host libraries were sequenced on NextSeq 550 (Illumina) using 1 x 75bp read length. Sequenced reads were aligned to human genome version GRCh38 p13 and quantified using the zUMIs pipeline and default parameters.^102^

### Statistical analysis of gene expression

Differential expression analysis was performed on raw counts using DESeq.^103^ For host and pathogen, log_2_ fold changes were moderated using apeGLM,^46^ and genes were considered significantly differentially expressed using an adjusted p value threshold < 0.05 and post-hoc filter of |moderated log_2_ fold change| > 1. To identify genes are not significantly differentially expressed under a condition, we performed a Wald test on unshrunk fold changes using the null hypothesis |log 2 fold changes| > 0.7 and an adjusted p value threshold < 0.05 (Supplementary Figure 8).

Enrichment of Mtb gene modules^50^ in shared and drug-specific sets of differentially expressed genes was calculated by hypergeometric test against the background set of all differentially expressed genes (n=320). Gene set enrichment analysis of the sigE regulon was performed on ordered, shrunk log2 fold changes using clusterProfiler’s implementation of GSEA, default parameters, and the sigE regulon defined by Manganelli et al,^47^ the marR regulon defined by Golby et al,^104^ and the kstR regulon defined by Kendall et al.^58^

## Code availability

All data, code, and instructions needed to reproduce computational analyses is available on https://github.com/cgunnars/egfr-seq_repo.

## Acknowledgments

Mtb work was performed in the Ragon Institute BSL3 core facility, which is supported by the NIH-funded Harvard University Center for AIDS Research (P30 AI060354). We thank Amy Barczak, Yong Xie, Julie Boucau, Eliane Shwiari, and Stephanie Pringle for managing the facility and advising on validation of new protocols for Mtb inactivation. Stuart Levine in the MIT BioMicrocenter assisted with design of Mtb-selective RNA probes, and BioMicrocenter student workers quantitated the sequencing libraries prior to loading. The White and Lauffenburger labs provided A549 cells. Ryan Milligan and Iris Abrahantes-Morales processed buffy coats. This work is supported by funding from NIH grants R35GM142900, R01AI184666 and R01A1022553 and the Army W911NF-19-D-0001.

## Supplementary Figure Captions

**Supplementary Figure 1.**
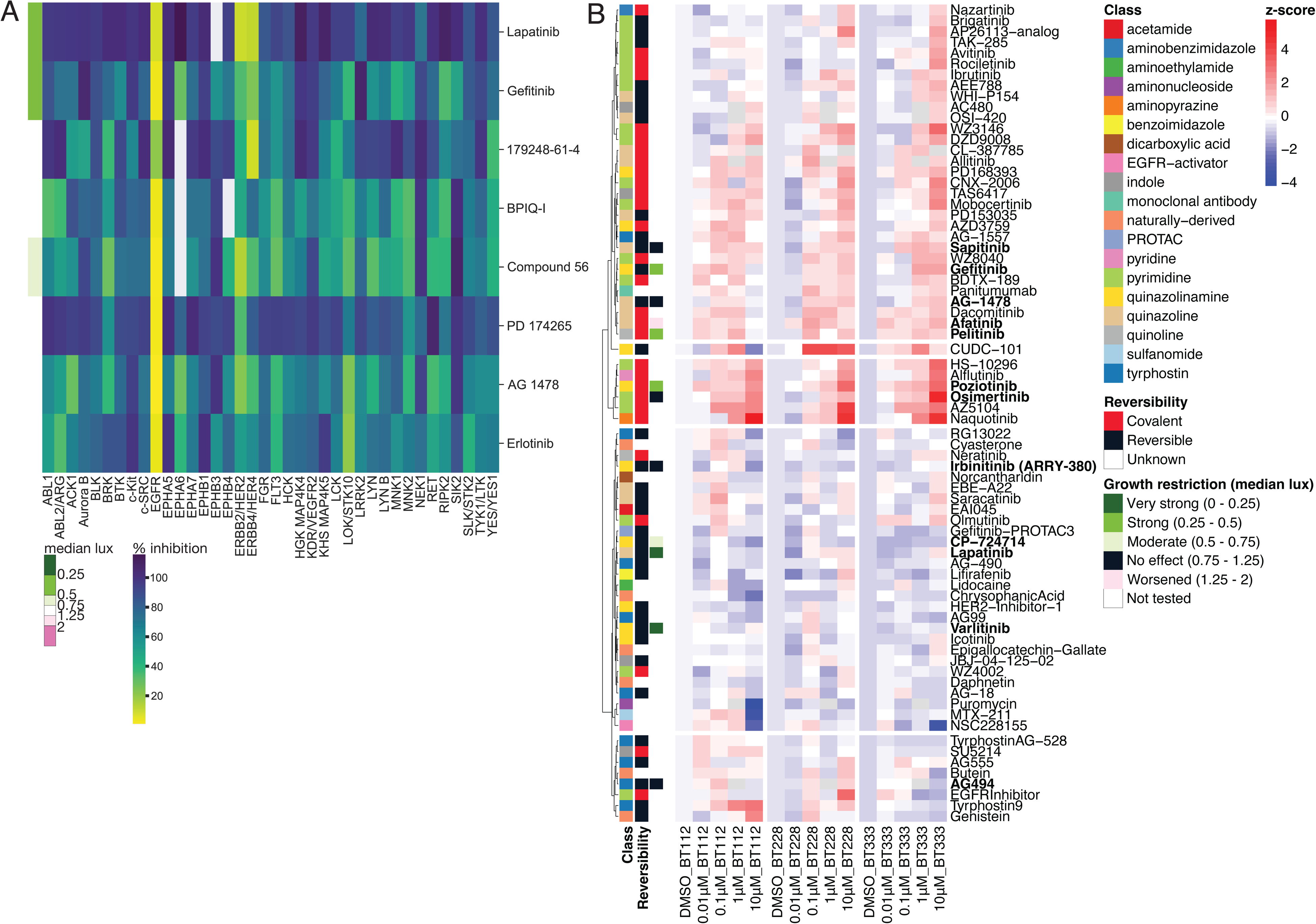
Known differences in inhibition of kinase catalytic activity or transcriptional response do not explain differences in intracellular growth restriction. Relates to Figure 1. **A)** Percent inhibition of kinase catalytic activity from Anastassiadis et al.^91^ is plotted for kinase targets with at least 75% inhibition by one drug. Compounds are annotated according to their effects on intracellular luminescence. **B)** Mean expression of EGFR inhibition response signature is plotted across a panel of EGFR inhibitors, three cell lines, and four doses. Data is reproduced from Giglio et al,^20^ and EGFR inhibitors tested for effect on intracellular Mtb are bolded and colored according to their effect on restriction (black=no effect, green=restrictive, pink=permissive).

**Supplementary Figure 2.**
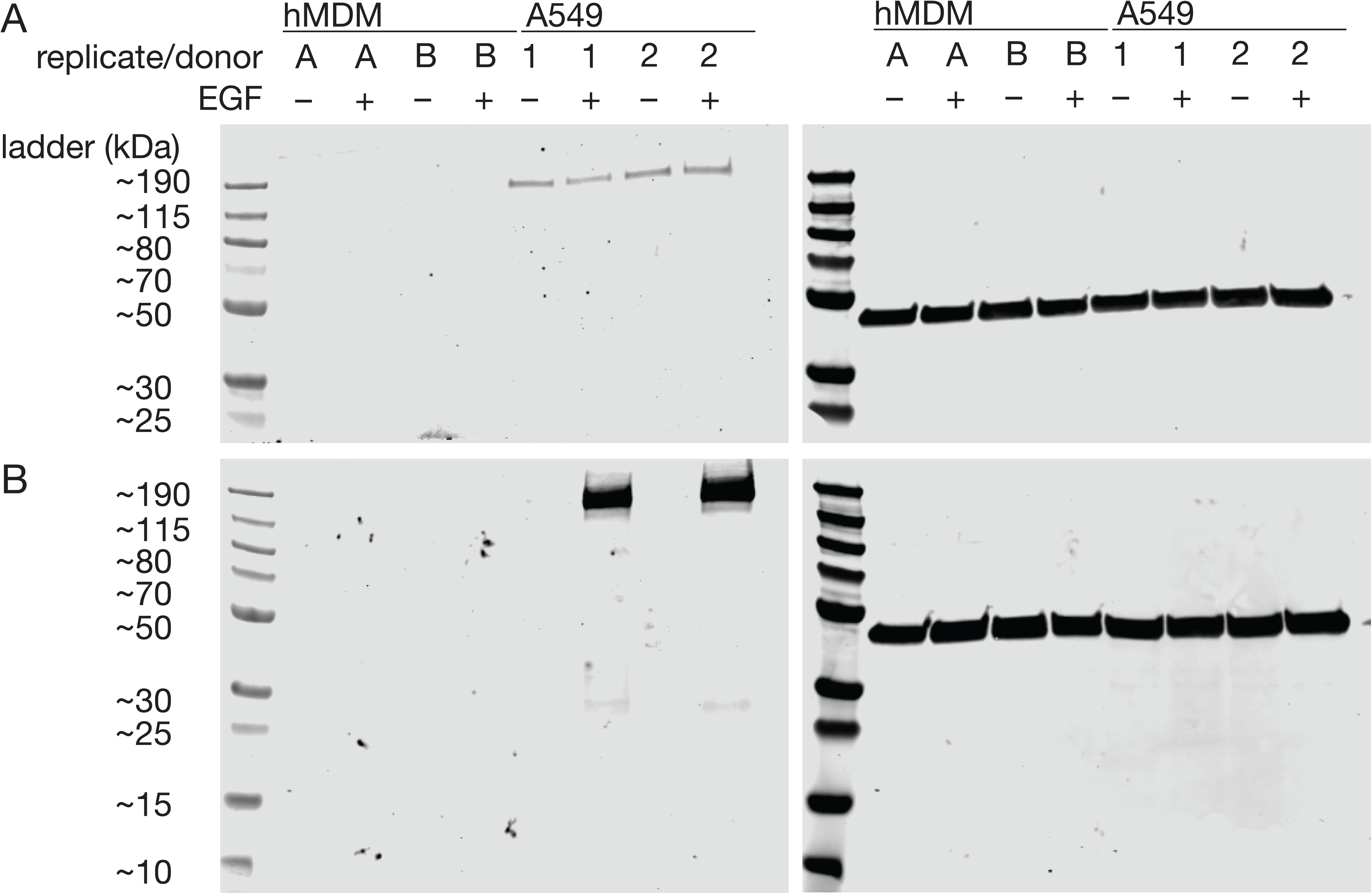
Human monocyte-derived macrophages express undetectable levels of total and phosphorylated EGFR. **Relates to Figure 1 A - B)** Total protein was isolated from human monocyte-derived macrophages or A549 cells, stimulated with 100 ng/mL recombinant EGF for 15 minutes, and western blotting was performed for total EGFR (**A**) and phospho-EGFR (**B**). Apparent molecular weight is annotated according to estimates for PageRuler Plus Prestained Protein Ladder, 4-12% Bis-Tris gel, MES running buffer. Beta-actin is shown as a loading control (right). Shown for n=2 donors for hMDM, 2 replicate wells for A549; representative of n=4 total donors isolated in two independent experiments.

**Supplementary Figure 3.**
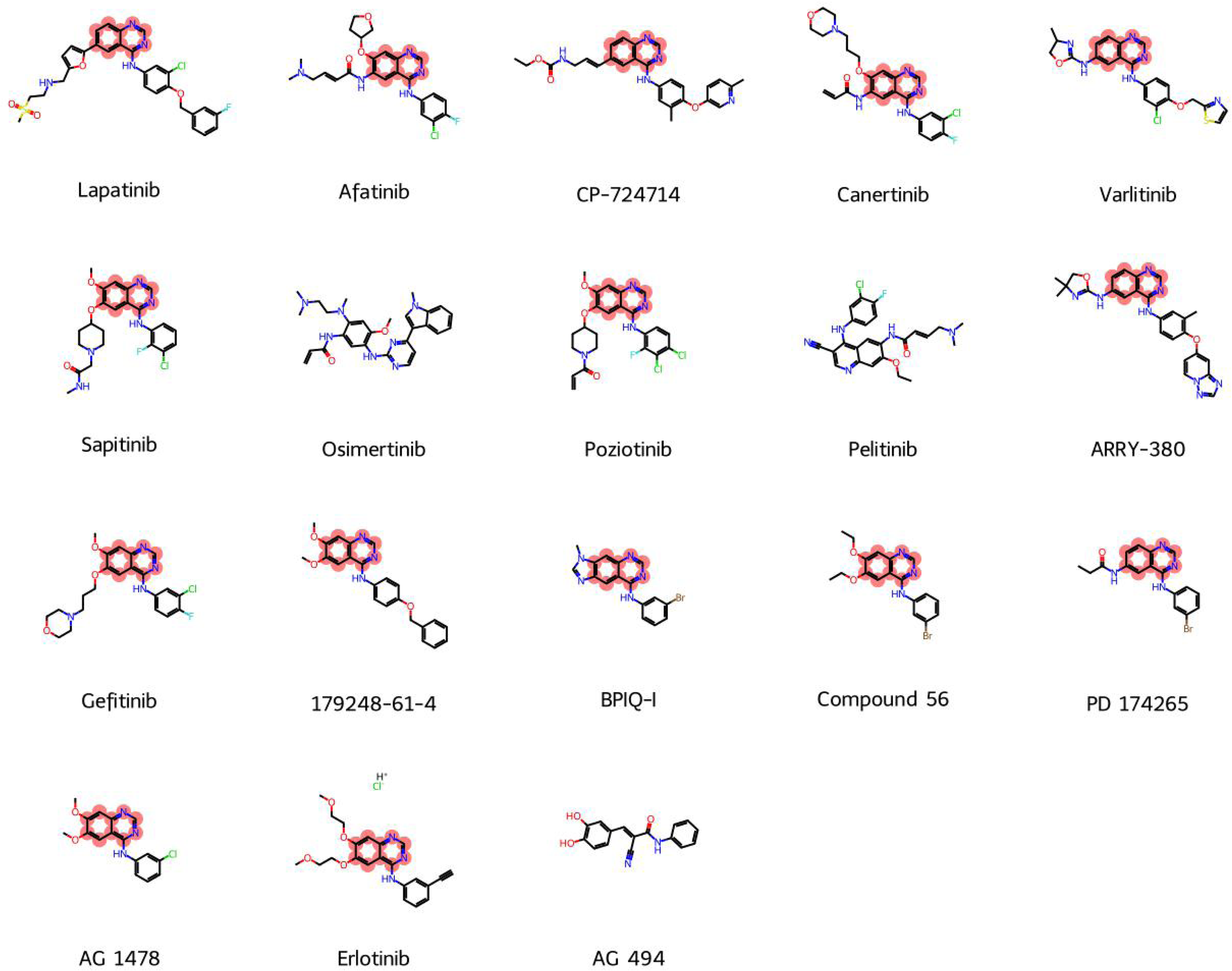
Chemical structures of EGFR inhibitors in panel. Relates to. Figure 2 Quinazoline scaffold is highlighted in red.

**Supplementary Figure 4.**
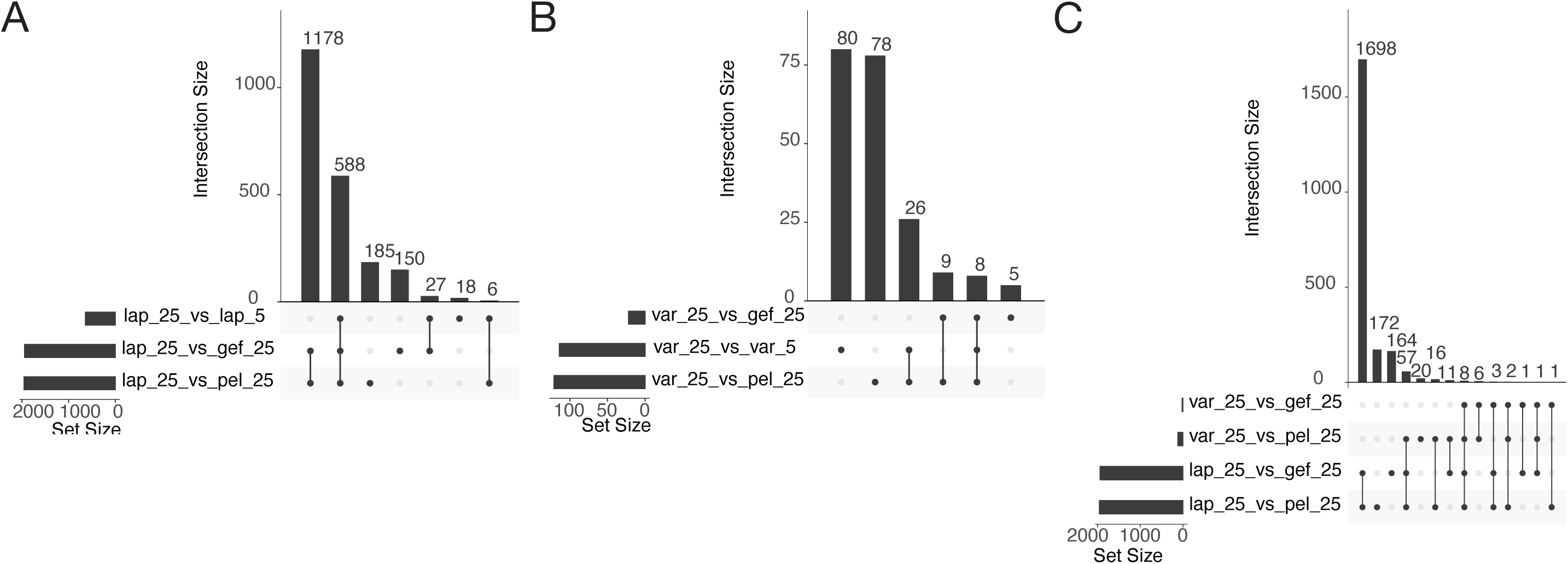
Overlap between varlitinib- and lapatinib-induced genes. **Relates to Figure 2 A - C)** Differential gene expression of Mtb treated with 25 µM lapatinib or varlitinib was calculated relative to 25 µM gefitinib/pelitinib or 5 µM lapatinib/varlitinib. The number and overlap in differentially expressed genes between comparisons is shown for high-dose lapatinib relative to non-restrictive references (**A**), high-dose varlitinib relative to non-restrictive references, (**B**), and high-dose lapatinib and high-dose varlitinib jointly (**C**).

**Supplementary Figure 5.**
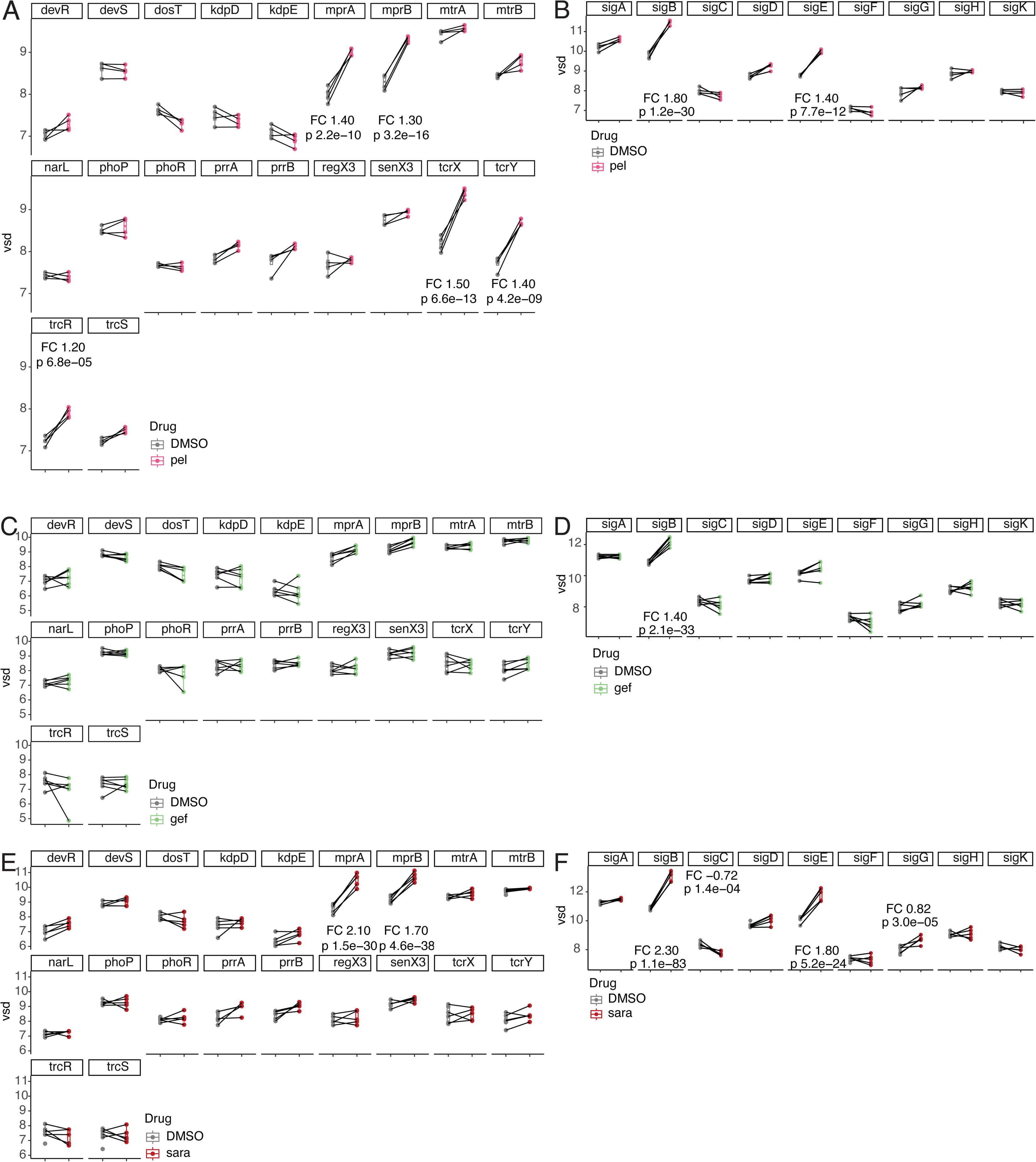
Treatment of infected macrophages with host-dependent therapy transcriptionally upregulates environment-responsive sensors and regulators. **Relates to Figure 4 A - F)** Transcript expression of intracellular Mtb was measured at 24 hours post-infection and treatment with 10 µM pelitinib, gefitinib, saracatinib, or vehicle. Variance-stabilized counts of Mtb two-component systems (**A, C, E**) and sigma factors (**B, D, F**) are plotted relative to vehicle for pelitinib (**A, B**), gefitinib (**C, D**), and saracatinib (**E, F**). For transcripts with |moderated log_2_ fold changes > 0.7|, moderated log_2_ fold changes and Benjamini-Hochberg–adjusted p values are listed. Points represent n=5-6 donors per condition.

**Supplementary Figure 6.**
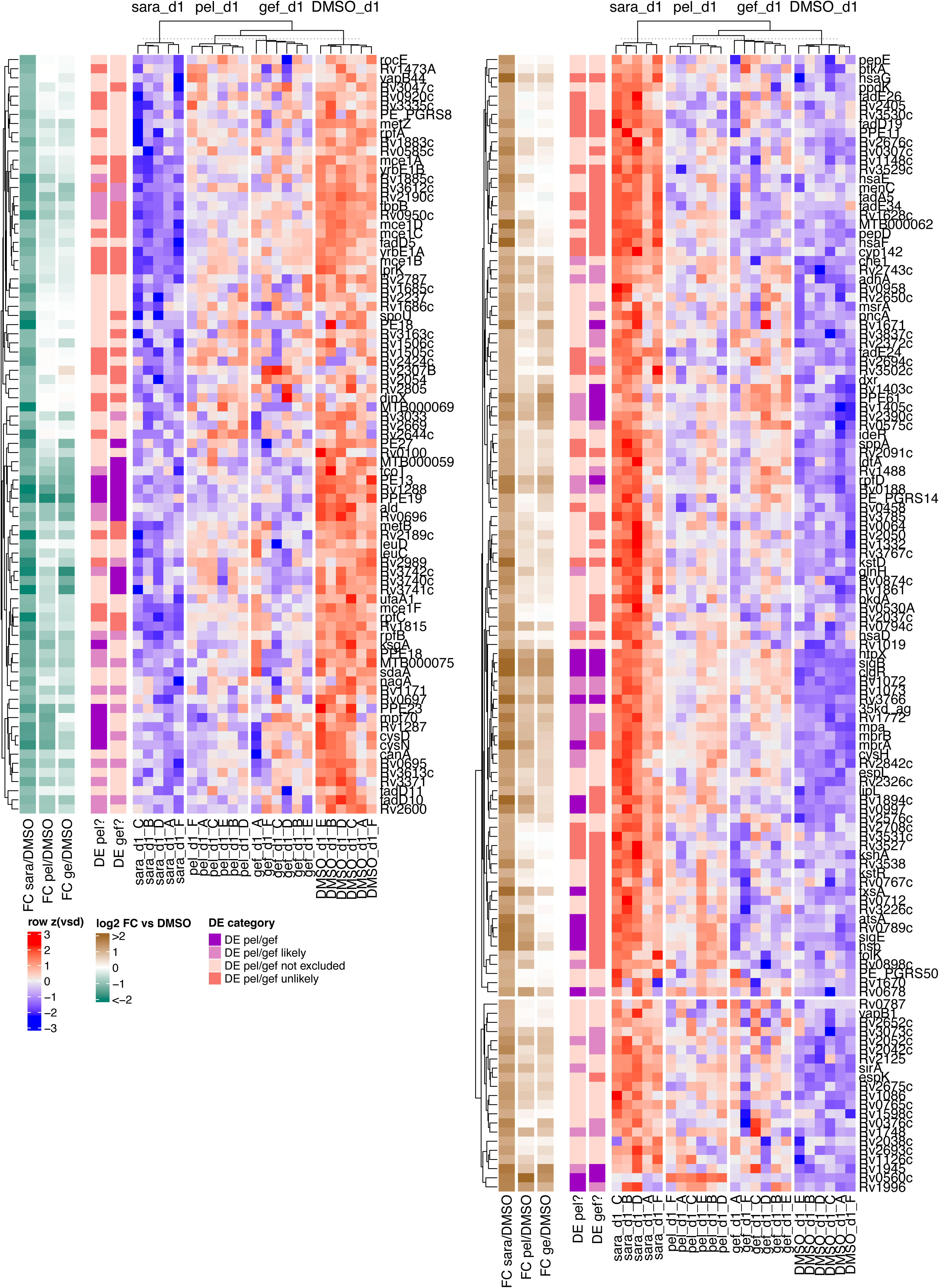
Mtb differential expression in saracatinib-treated macrophages. Relates to. Figure 4 Differentially expressed genes in the saracatinib condition relative to the DMSO condition. For each gene, variance-stabilized counts at 24 hours post-infection are z-scored and plotted across treatment conditions. Moderated estimates of per-gene log_2_ fold change are also plotted for each condition relative to DMSO, and each gene is statistically categorized according to whether it is differentially expressed under non-saracatinib conditions relative to DMSO.

**Supplementary Figure 7.**
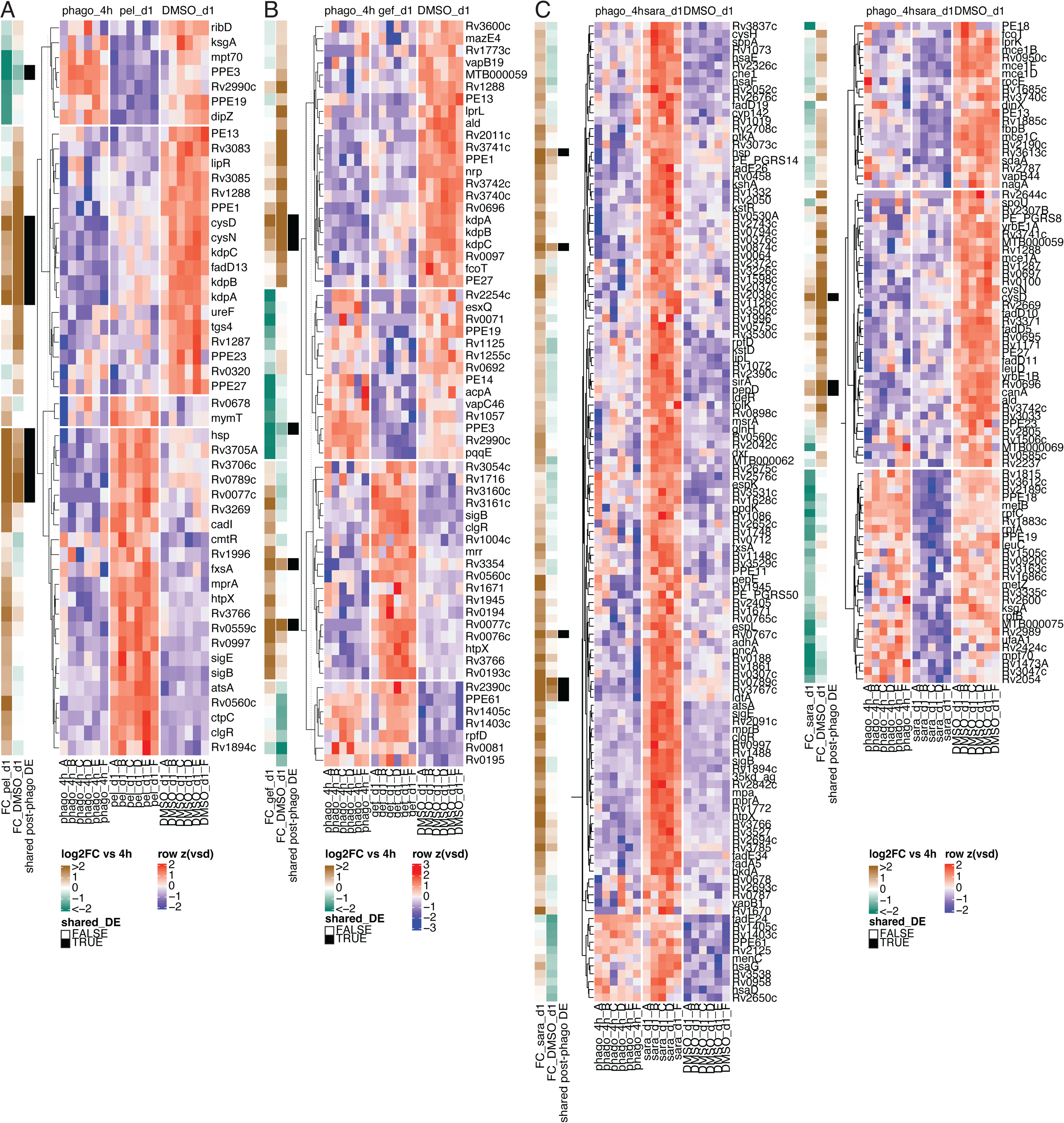
Host-dependent therapy changes the trajectory of post-phagocytosis induction. **Relates to Figure 4 A-C)** Expression of significantly differentially expressed genes between pelitinib-(**A**), gefitinib-(**B**), or saracatinib-(**C**) and vehicle-treated macrophages at 24 hours is plotted for samples isolated post-phagocytosis, but pre-treatment and samples isolated at 24 hours.

**Supplementary Figure 8.**
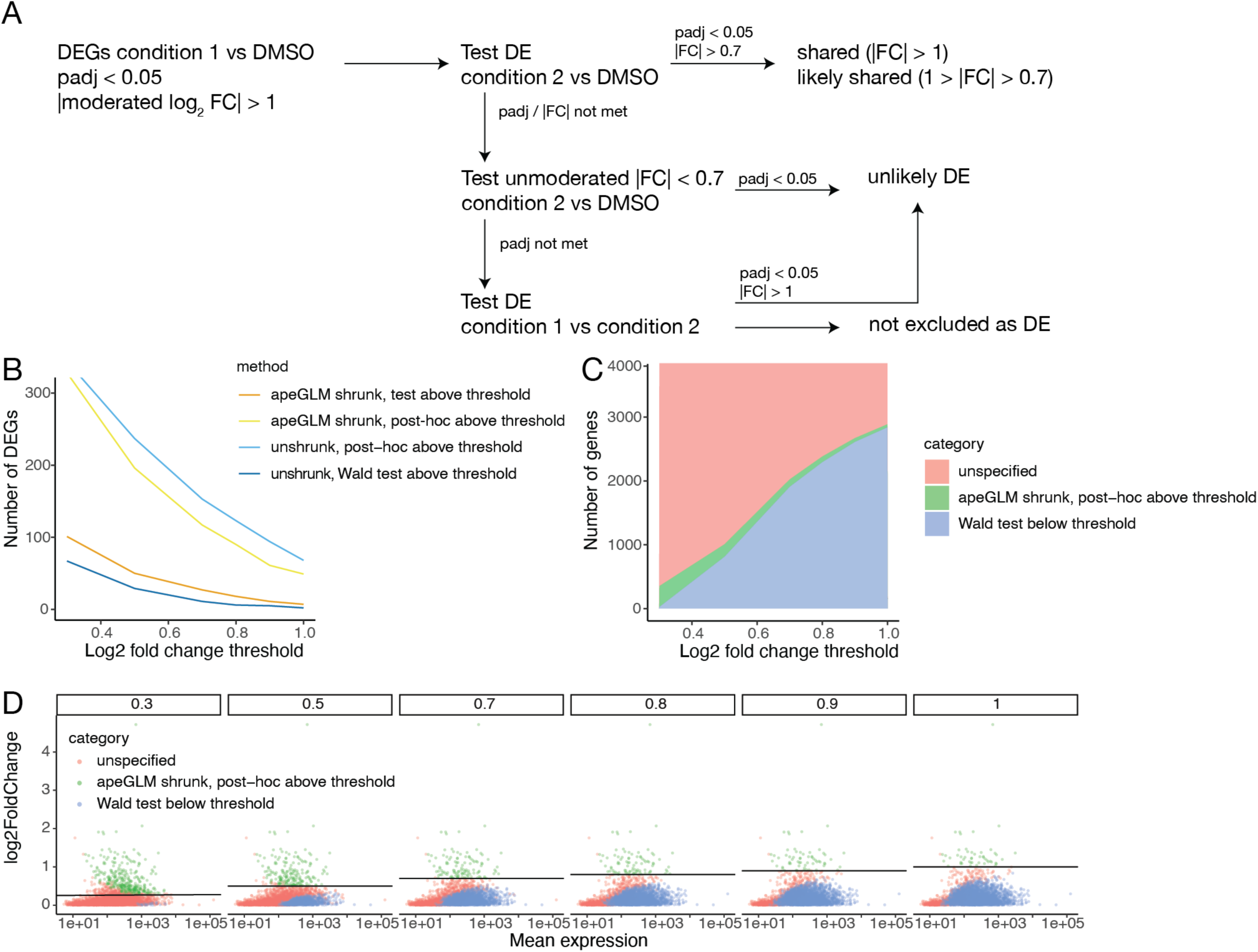
Statistical framework for defining low-change genes. Relates to. **Figure 4 A)** Overview of statistical approach. **B)** Benchmarking of four differential expression methods. The number of differentially expressed genes is plotted against a range of log_2_ fold changes. For all tests, a Benjamini-Hochberg–adjusted p value threshold of 0.05 was used. **C)** Effect of log_2_ fold change threshold on determination of differentially expressed genes, low-change genes, and genes unspecified by either test. **D)** Effect of mean expression on gene categorization as a function of log_2_ fold change thresholds used in statistical testing. For each subpanel, a horizontal line corresponding to the log_2_ fold change threshold is plotted.

**Supplementary Figure 9.**
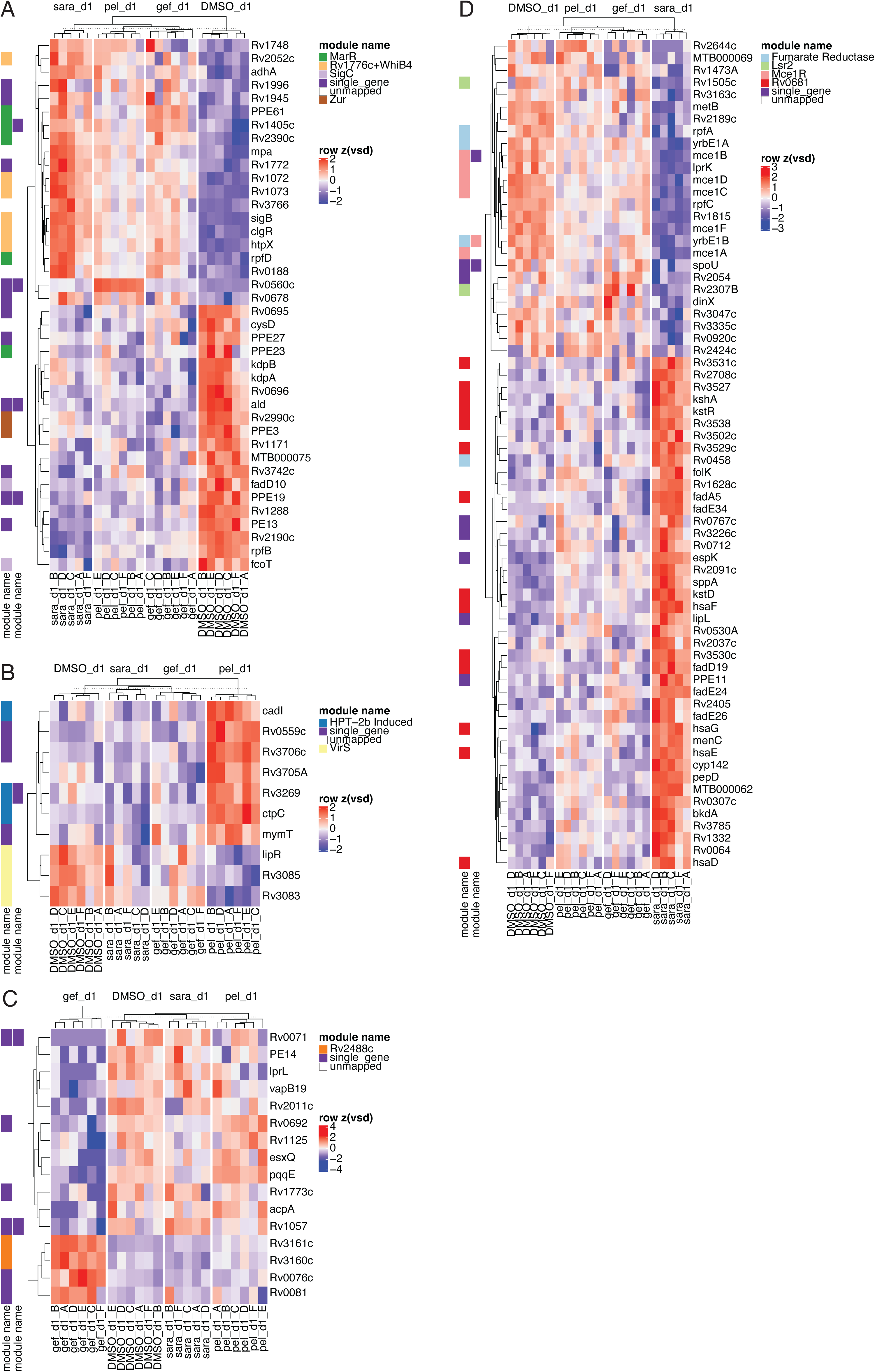
Shared and drug-specific Mtb genes with differential expression relative to vehicle. **Relates to Figure 4 A - D)** Differentially expressed genes for treatment conditions relative to vehicle at 24 hours are defined as shared across all treatment conditions (**A**), pelitinib-specific (**B**), gefitinib-specific (**C**) or saracatinib-specific (**D**). For each gene, variance-stabilized counts at 24 hours post-infection are z-scored and annotated according to mapping to iModulon-defined transcriptional modules.^50^

**Supplementary Figure 10.**
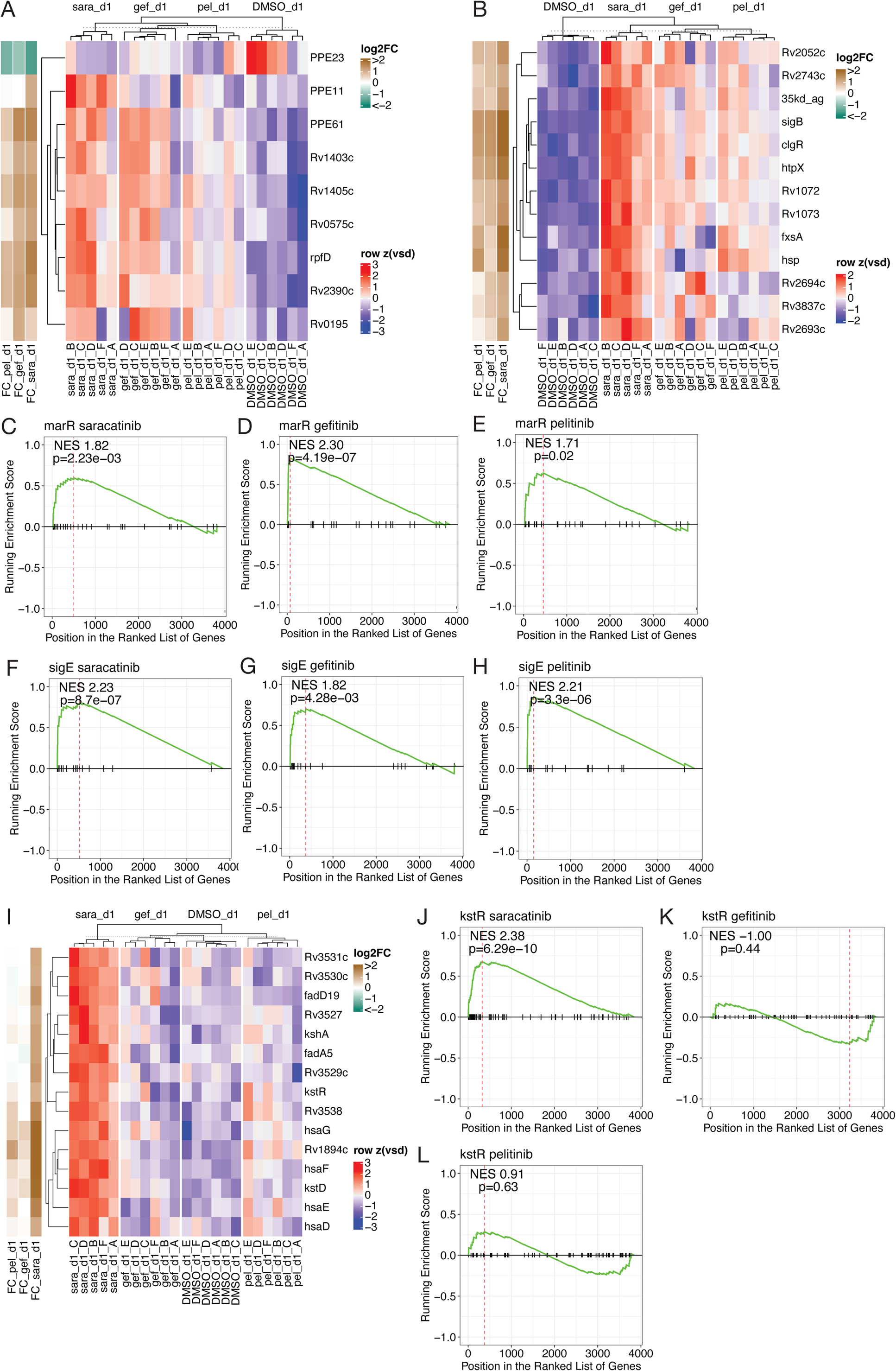
Module enrichment in shared and drug-specific gene lists is supported by expression patterns and orthogonal enrichment analysis. **Relates to Figure 4 A - B)** Variance-stabilized counts of genes that contributed to shared enrichment of **(A)** MarR and **(B)** Rv1776c+WhiB4 iModulons are z-scored and plotted across all treatment conditions. **C - H).** Gene set enrichment analysis of the marR (**C-E**) and sigE (**F-H**) regulons was conducted on ranked, moderated log2 fold change estimates for each condition relative to vehicle. Normalized enrichment scores and adjusted p-values are shown on each plot. **I)** Variance-stabilized counts of genes that contributed to shared enrichment of the Rv0681 iModulon is z-scored and plotted across all treatment conditions. **J - L)** Gene set enrichment analysis of the kstR regulon for saracatinib (**J**), gefitinib (**K**) and pelitinib DEGs (**L**). Multiple hypothesis correction was performed using the Benjamini-Hochberg correction for each condition separately, considering each gene set as a separate hypothesis.

**Supplementary Figure 11.**
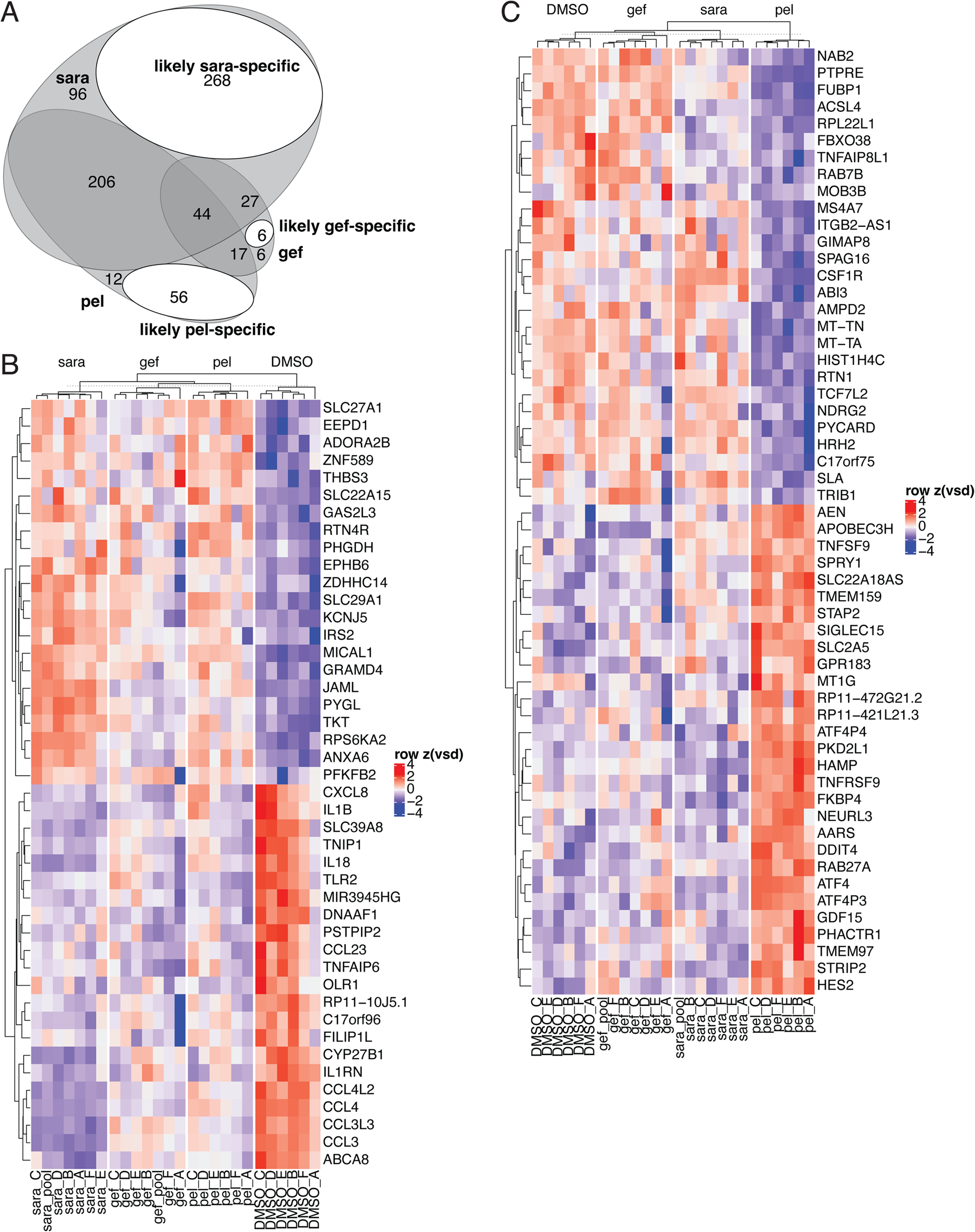
Host transcriptional state in Mtb-infected macrophages treated with host-dependent therapy. **Relates to Figure 4 A)** Paired host RNA-seq libraries were generated from RNA isolated in the experiment described by Figure 4B for n=5-6 donors per condition. Differentially expressed host genes (|moderated log_2_ FC| > 1, padj < 0.05) at 24 hours post-infection was calculated for each drug condition relative to DMSO. Overlap in likely DEGs relative to DMSO across pelitinib, gefitinib, and saracatinib conditions. Likely drug-specific genes within non-overlapping regions of the Venn diagram are highlighted. **B)** Expression of likely shared host DEGs across all treatment conditions. For each gene, variance-stabilized counts at 24 hours post-infection are z-scored and plotted across treatment conditions. **C)** Expression of pelitinib-specific host DEGs.

**Supplementary Figure 12.**
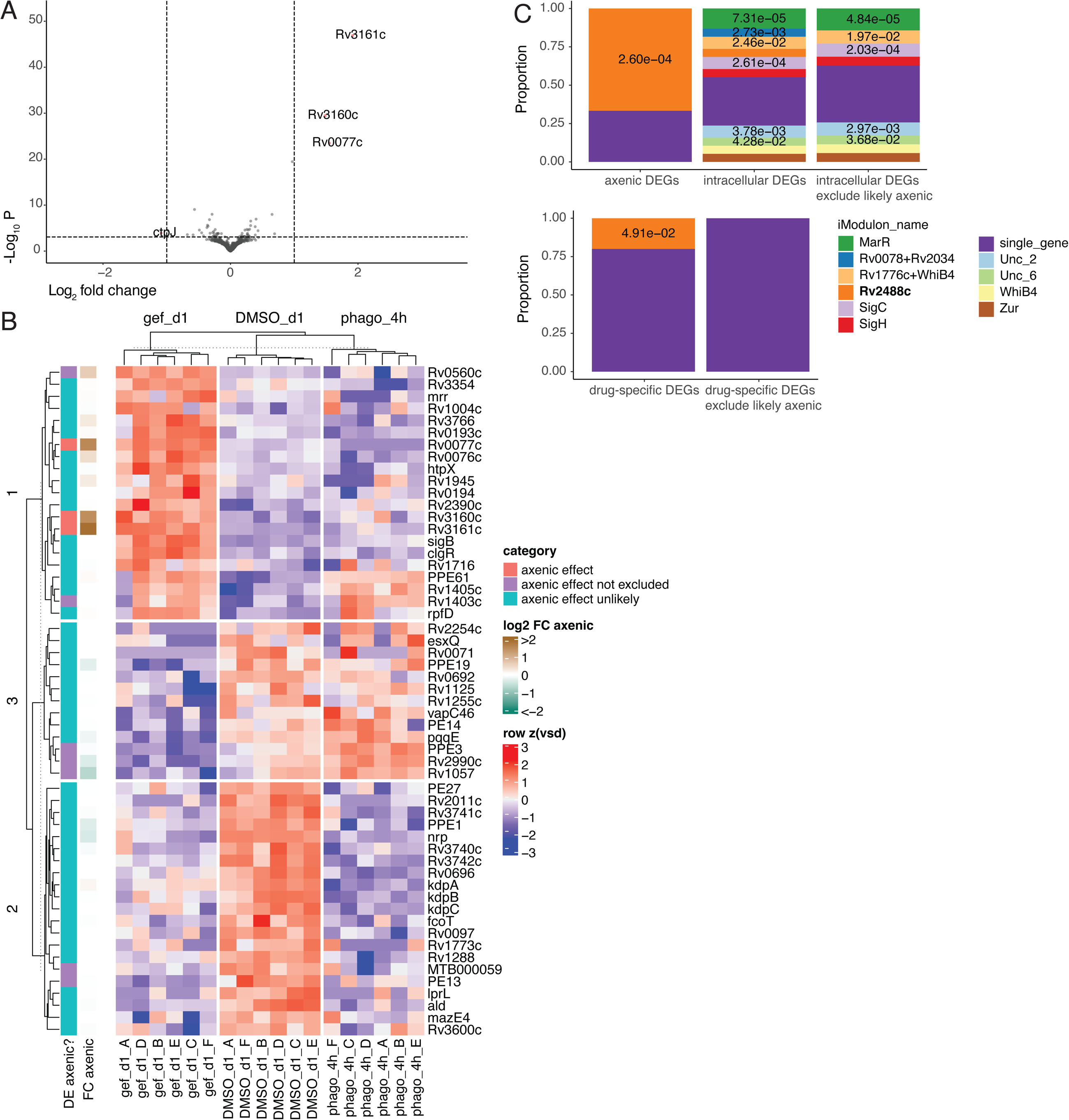
Effects of gefitinib therapy on Mtb gene expression include host environment-dependent and axenic effects. **Relates to Figure 5 A)** Differential expression was calculated for axenic Mtb cultures exposed to 10 µM gefitinib or vehicle in 7H9 for 24 hours, n=4 biological replicates. Log_10_ Benjamini-Hochberg–adjusted p values are plotted against moderated log_2_ fold change estimates for each gene, and genes with significant differential expression are colored in red. **B)** Heatmap of intracellular differential expression in gefitinib relative to DMSO at 24 hours, annotated according to moderated log_2_ fold change in axenic conditions and whether genes are statistically categorized as exhibiting axenic differential expression. **C)** Hypergeometric enrichment of iModulonDB-defined Mtb gene modules^50^ in gefitinib and gefitinib-specific DEGs, excluding genes with axenic differential expression. For gefitinib DEGs, the background set is defined as all iModulon-mappable genes, while for gefitinib-specific DEGs, the background set is defined as all DEGs (n = 320).

**Supplementary Table 1.** Shared genes with dose-dependent differential expression in axenic Mtb. For each condition, genes with significant differential expression (moderated |log2FC| > 1; padj < 0.05) are considered.

|  | N genes | Gene names |
| --- | --- | --- |
| Shared (all) | 7 | Downregulated: cyp121, rpfE, PPE18 (upregulated by high-dose lapatinib)<br>Upregulated: ctpJ, Rv3160c, Rv0678, mmpL5 |
| Pelitinib +<br>gefitinib +<br>varlitinib | 8 | Downregulated: MTB000075, PE13, esxK, esxL, modA<br>Upregulated: prpD, prpC, frdB |
| Pelitinib +<br>gefitinib +<br>lapatinib | 3 | Downregulated: lipR (upregulated by high-dose lapatinib)<br>Upregulated: mprA, Rv2275 |
| Pelitinib +<br>lapatinib +<br>varlitinib | 4 | Downregulated: Rv0888, Rv3083 (upregulated by high-dose lapatinib)<br>Upregulated: sigE, Rv0560c |
| Gefitinib +<br>lapatinib +<br>varlitinib | 6 | Downregulated: PPE3<br>Upregulated: Rv3161c, Rv3269, ctpC, lat, PPE53 |

**Supplementary Table 2.** Stringently shared differentially expressed genes in intracellular Mtb. For each condition, genes with significant differential expression above vehicle (moderated |log2FC| > 1; padj < 0.05) are considered.

|  | N genes | Gene names |
| --- | --- | --- |
| Shared (all) | 8 | Downregulated: PE13, Rv1288, PPE19<br>Upregulated: clgR, htpX, sigB, Rv3766, Rv0560c |
| Pelitinib + gefitinib | 7 | Downregulated: PPE1, PPE3, Rv2990c, kdpA, kdpB, kdpC<br>Upregulated: Rv0077c |
| Pelitinib + saracatinib | 16 | Downregulated: ksgA, Rv1287, cysD, cysN, PPE23, mpt70<br>Upregulated: fxsA, Rv0997, Rv1984c, hsp, mprA, atsA, Rv0789c, sigE, Rv1996, Rv0678 |
| Gefitinib + saracatinib | 15 | Downregulated: MTB000059, fcoT, PE27, Rv3740c, Rv3741c, Rv3742c, ald, Rv0696<br>Upregulated: rpfD, Rv1403c, Rv1405c, PPE61, Rv1671, Rv1945, Rv2390c |

## Tables

Supplementary Table 3. Differential expression results for axenic Mtb treated with host-independent drugs at 25 µM for 4h, relative to host-dependent drugs

Supplementary Table 4. Dose-dependent differential expression results for axenic Mtb treated with drug at 25 µM vs 5 µM for 4h

Supplementary Table 5. Differential expression results for intracellular Mtb treated with host-dependent drugs at 10µM for 24h, relative to DMSO at 24h

Supplementary Table 6. Differential expression results for axenic Mtb treated with host-dependent drugs at 10µM for 24h, relative to DMSO at 24h

Supplementary Table 7. Hypergeometric iModulon enrichment results for shared and drug-specific DEGs

Supplementary Table 8. Gene set enrichment results for intracellular DEGs

Supplementary Table 9. Hypergeometric iModulon enrichment results for intracellular pelitinib and pelitinib-specific DEGs, with and without exclusion of genes with axenic differential expression

Supplementary Table 10. Hypergeometric iModulon enrichment results for intracellular pelitinib and gefitinib-specific DEGs, with and without exclusion of genes with axenic differential expression

## References

1. Connolly, L. E., Edelstein, P. H. & Ramakrishnan, L. Why is long-term therapy required to cure tuberculosis? PLoS Med. (2007) doi:10.1371/journal.pmed.0040120.

2. Kaufmann, S. H. E., Dorhoi, A., Hotchkiss, R. S. & Bartenschlager, R. Host-directed therapies for bacterial and viral infections. Nat. Rev. Drug Discov. 17, 35–56 (2018).

3. Shapira, T. et al. High-Content Screening of Eukaryotic Kinase Inhibitors Identify CHK2 Inhibitor Activity Against Mycobacterium tuberculosis. Front. Microbiol. (2020) doi:10.3389/fmicb.2020.553962.

4. Korbee, C. J. et al. Combined chemical genetics and data-driven bioinformatics approach identifies receptor tyrosine kinase inhibitors as host-directed antimicrobials. Nat. Commun. (2018) doi:10.1038/s41467-017-02777-6.

5. Stanley, S. A. et al. Identification of Host-Targeted Small Molecules That Restrict Intracellular Mycobacterium tuberculosis Growth. PLoS Pathog. (2014) doi:10.1371/journal.ppat.1003946.

6. Hardbower, D. M. et al. EGFR regulates macrophage activation and function in bacterial infection. J. Clin. Invest. 126, 3296–3312 (2016).

7. Zhang, W. et al. Polarization of macrophages in the tumor microenvironment is influenced by EGFR signaling within colon cancer cells. Oncotarget 7, 75366–75378 (2016).

8. Lu, N. et al. Activation of the Epidermal Growth Factor Receptor in Macrophages Regulates Cytokine Production and Experimental Colitis. J. Immunol. 192, 1013–1023 (2014).

9. van den Biggelaar, R. H. G. A., et al. Identification of kinase inhibitors as potential host-directed therapies for intracellular bacteria. Sci. Rep. 14, 17225 (2024).

10. Sadhu, S. et al. Gefitinib Results in Robust Host-Directed Immunity Against Salmonella Infection Through Proteo-Metabolomic Reprogramming. Front. Immunol. 12, 648710 (2021).

11. van den Biggelaar, R. H. G. A., et al. Identification of kinase modulators as host-directed therapeutics against intracellular methicillin-resistant Staphylococcus aureus. Front. Cell. Infect. Microbiol. 14, 1367938 (2024).

12. Lai, Y. et al. Illuminating Host-Mycobacterial Interactions with Genome-wide CRISPR Knockout and CRISPRi Screens. Cell Syst. 11, 239–251.e7 (2020).

13. Sogi, K. M., Lien, K. A., Johnson, J. R., Krogan, N. J. & Stanley, S. A. The Tyrosine Kinase Inhibitor Gefitinib Restricts Mycobacterium tuberculosis Growth through Increased Lysosomal Biogenesis and Modulation of Cytokine Signaling. ACS Infect. Dis. 3, 564–574 (2017).

14. de Chastellier, C., Forquet, F., Gordon, A. & Thilo, L. Mycobacterium requires an all-around closely apposing phagosome membrane to maintain the maturation block and this apposition is re-established when it rescues itself from phagolysosomes. Cell. Microbiol. 11, 1190–1207 (2009).

15. Phagosome-lysosome interactions in cultured macrophages infected with virulent tubercle bacilli. Reversal of the usual nonfusion pattern and observations on bacterial survival. J. Exp. Med. 142, 1–16 (1975).

16. Andreu, N., et al. Optimisation of Bioluminescent Reporters for Use with Mycobacteria. PLOS ONE 5, e10777 (2010).

17. Andreu, N., Fletcher, T., Krishnan, N., Wiles, S. & Robertson, B. D. Rapid measurement of antituberculosis drug activity in vitro and in macrophages using bioluminescence. J. Antimicrob. Chemother. 67, 404–414 (2012).

18. Andreu, N. et al. Rapid in vivo assessment of drug efficacy against Mycobacterium tuberculosis using an improved firefly luciferase. J. Antimicrob. Chemother. 68, 2118–2127 (2013).

19. Blumenberg, M. Differential Transcriptional Effects of EGFR Inhibitors. PLOS ONE 9, e102466 (2014).

20. Giglio, R. M. et al. A heterogeneous pharmaco-transcriptomic landscape induced by targeting a single oncogenic kinase. BioRxiv Prepr. Serv. Biol. 2024.04.08.587960 (2025) doi:10.1101/2024.04.08.587960.

21. Dutta, A. & Sarma, D. Recent advances in the synthesis of Quinazoline analogues as Anti-TB agents. Tuberc. Edinb. Scotl. 124, 101986 (2020).

22. Asquith, C. R. M. et al. Anti-tubercular activity of novel 4-anilinoquinolines and 4-anilinoquinazolines. Bioorg. Med. Chem. Lett. 29, 2695–2699 (2019).

23. Franzblau, S. G. et al. Rapid, Low-Technology MIC Determination with Clinical Mycobacterium tuberculosis Isolates by Using the Microplate Alamar Blue Assay. J. Clin. Microbiol. 36, 362–366 (1998).

24. Collins, L. & Franzblau, S. G. Microplate alamar blue assay versus BACTEC 460 system for high-throughput screening of compounds against Mycobacterium tuberculosis and Mycobacterium avium. Antimicrob. Agents Chemother. 41, 1004–1009 (1997).

25. Nadanaciva, S. et al. A high content screening assay for identifying lysosomotropic compounds. Toxicol. Vitro Int. J. Publ. Assoc. BIBRA 25, 715–723 (2011).

26. Smollett, K. L. et al. Global analysis of the regulon of the transcriptional repressor LexA, a key component of SOS response in Mycobacterium tuberculosis. J. Biol. Chem. 287, 22004–22014 (2012).

27. Boshoff, H. I. M. et al. The transcriptional responses of Mycobacterium tuberculosis to inhibitors of metabolism: novel insights into drug mechanisms of action. J. Biol. Chem. 279, 40174–40184 (2004).

28. Shee, S. et al. Moxifloxacin-Mediated Killing of Mycobacterium tuberculosis Involves Respiratory Downshift, Reductive Stress, and Accumulation of Reactive Oxygen Species. Antimicrob. Agents Chemother. 66, e0059222 (2022).

29. Kim, N.-K., Baek, J.-E., Lee, Y.-J., Oh, Y. & Oh, J.-I. Rel-dependent decrease in the expression of ribosomal protein genes by inhibition of the respiratory electron transport chain in Mycobacterium smegmatis. Front. Microbiol. 15, 1448277 (2024).

30. Iacobino, A., Piccaro, G., Pardini, M., Fattorini, L. & Giannoni, F. Moxifloxacin Activates the SOS Response in Mycobacterium tuberculosis in a Dose- and Time-Dependent Manner. Microorganisms 9, 255 (2021).

31. Betts, J. C., Lukey, P. T., Robb, L. C., McAdam, R. A. & Duncan, K. Evaluation of a nutrient starvation model of Mycobacterium tuberculosis persistence by gene and protein expression profiling. Mol. Microbiol. 43, 717–731 (2002).

32. Keren, I., Minami, S., Rubin, E. & Lewis, K. Characterization and Transcriptome Analysis of Mycobacterium tuberculosis Persisters. mBio 2, 10.1128/mbio.00100-11 (2011).

33. Hasenoehrl, E. J. et al. Derailing the aspartate pathway of Mycobacterium tuberculosis to eradicate persistent infection. Nat. Commun. 10, 4215 (2019).

34. Duan, X. et al. Mycobacterium Lysine ε-aminotransferase is a novel alarmone metabolism related persister gene via dysregulating the intracellular amino acid level. Sci. Rep. 6, 19695 (2016).

35. Liu, Y. et al. NapM enhances the survival of Mycobacterium tuberculosis under stress and in macrophages. *Commun*. Biol. 2, 65 (2019).

36. Aguilar-Ayala, D. A. et al. The transcriptome of Mycobacterium tuberculosis in a lipid-rich dormancy model through RNAseq analysis. Sci. Rep. 7, 17665 (2017).

37. Watanabe, S. et al. Fumarate Reductase Activity Maintains an Energized Membrane in Anaerobic Mycobacterium tuberculosis. PLOS Pathog. 7, e1002287 (2011).

38. Mavi, P. S., Singh, S. & Kumar, A. Reductive Stress: New Insights in Physiology and Drug Tolerance of Mycobacterium. Antioxid. Redox Signal. 32, 1348–1366 (2020).

39. Small, J. L. et al. Perturbation of cytochrome c maturation reveals adaptability of the respiratory chain in Mycobacterium tuberculosis. mBio 4, e00475–00413 (2013).

40. Cai, Y. et al. Host immunity increases Mycobacterium tuberculosis reliance on cytochrome bd oxidase. PLOS Pathog. 17, e1008911 (2021).

41. Schnappinger, D. et al. Transcriptional Adaptation of Mycobacterium tuberculosis within Macrophages. J. Exp. Med. 198, 693–704 (2003).

42. Pisu, D., Huang, L., Grenier, J. K. & Russell, D. G. Dual RNA-Seq of Mtb-Infected Macrophages In Vivo Reveals Ontologically Distinct Host-Pathogen Interactions. Cell Rep. (2020) doi:10.1016/j.celrep.2019.12.033.

43. Torrance, C. J. et al. Combinatorial chemoprevention of intestinal neoplasia. Nat. Med. 6, 1024–1028 (2000).

44. Healy, C., Golby, P., MacHugh, D. E. & Gordon, S. V. The MarR family transcription factor Rv1404 coordinates adaptation of Mycobacterium tuberculosis to acid stress via controlled expression of Rv1405c, a virulence-associated methyltransferase. Tuberc. Edinb. Scotl. 97, 154–162 (2016).

45. MacGilvary, N. J., Kevorkian, Y. L. & Tan, S. Potassium response and homeostasis in Mycobacterium tuberculosis modulates environmental adaptation and is important for host colonization. PLoS Pathog. 15, e1007591 (2019).

46. Zhu, A., Ibrahim, J. G. & Love, M. I. Heavy-tailed prior distributions for sequence count data: removing the noise and preserving large differences. Bioinformatics 35, 2084–2092 (2019).

47. Manganelli, R., Voskuil, M. I., Schoolnik, G. K. & Smith, I. The Mycobacterium tuberculosis ECF sigma factor sigmaE: role in global gene expression and survival in macrophages. Mol. Microbiol. 41, 423–437 (2001).

48. White, M. J., He, H., Penoske, R. M., Twining, S. S. & Zahrt, T. C. PepD Participates in the Mycobacterial Stress Response Mediated through MprAB and SigE. J. Bacteriol. 192, 1498–1510 (2010).

49. Stupar, M. et al. TcrXY is an acid-sensing two-component transcriptional regulator of Mycobacterium tuberculosis required for persistent infection. Nat. Commun. 15, 1615 (2024).

50. Yoo, R. et al. Machine Learning of All Mycobacterium tuberculosis H37Rv RNA-seq Data Reveals a Structured Interplay between Metabolism, Stress Response, and Infection. mSphere 7, e00033–22.

51. Salina, E. G. et al. Copper-related toxicity in replicating and dormant Mycobacterium tuberculosis caused by 1-hydroxy-5-R-pyridine-2(1H)-thiones. Met. Integr. Biometal Sci. 10, 992–1002 (2018).

52. Flentie, K. et al. Chemical disarming of isoniazid resistance in Mycobacterium tuberculosis. Proc. Natl. Acad. Sci. U. S. A. 116, 10510–10517 (2019).

53. Gomez, A., Andreu, N., Ferrer-Navarro, M., Yero, D. & Gibert, I. Triclosan-induced genes Rv1686c-Rv1687c and Rv3161c are not involved in triclosan resistance in Mycobacterium tuberculosis. Sci. Rep. 6, 26221 (2016).

54. Dutta, N. K., Mehra, S. & Kaushal, D. A Mycobacterium tuberculosis sigma factor network responds to cell-envelope damage by the promising anti-mycobacterial thioridazine. PloS One 5, e10069 (2010).

55. Tükenmez, H. et al. Mycobacterium tuberculosis Rv3160c is a TetR-like transcriptional repressor that regulates expression of the putative oxygenase Rv3161c. Sci. Rep. 11, 1523 (2021).

56. Nazarova, E. V. et al. Rv3723/LucA coordinates fatty acid and cholesterol uptake in Mycobacterium tuberculosis. eLife https://elifesciences.org/articles/26969 (2017) doi:10.7554/eLife.26969.

57. Van der Geize, R. et al. A gene cluster encoding cholesterol catabolism in a soil actinomycete provides insight into Mycobacterium tuberculosis survival in macrophages. Proc. Natl. Acad. Sci. 104, 1947–1952 (2007).

58. Kendall, S. L. et al. A highly conserved transcriptional repressor controls a large regulon involved in lipid degradation in Mycobacterium smegmatis and Mycobacterium tuberculosis. Mol. Microbiol. 65, 684–699 (2007).

59. Nelson, J. K. et al. EEPD1 Is a Novel LXR Target Gene in Macrophages Which Regulates ABCA1 Abundance and Cholesterol Efflux. Arterioscler. Thromb. Vasc. Biol. 37, 423–432 (2017).

60. Cubells, L. et al. Annexin A6-Induced Alterations in Cholesterol Transport and Caveolin Export from the Golgi Complex. Traffic Cph. Den. 8, 1568–1589 (2007).

61. Chui, P. C., Guan, H.-P., Lehrke, M. & Lazar, M. A. PPARγ regulates adipocyte cholesterol metabolism via oxidized LDL receptor 1. J. Clin. Invest. 115, 2244–2256 (2005).

62. Trigueros-Motos, L. et al. ABCA8 Regulates Cholesterol Efflux and High-Density Lipoprotein Cholesterol Levels. Arterioscler. Thromb. Vasc. Biol. 37, 2147–2155 (2017).

63. Wheelwright, M. et al. All-trans retinoic acid-triggered antimicrobial activity against Mycobacterium tuberculosis is dependent on NPC2. J. Immunol. Baltim. Md 1950 192, 2280–2290 (2014).

64. Bartlett, S. et al. GPR183 Regulates Interferons, Autophagy, and Bacterial Growth During Mycobacterium tuberculosis Infection and Is Associated With TB Disease Severity. Front. Immunol. 11, 601534 (2020).

65. Sow, F. B. et al. Expression and localization of hepcidin in macrophages: a role in host defense against tuberculosis. J. Leukoc. Biol. 82, 934–945 (2007).

66. Maure, A. et al. A host-directed oxadiazole compound potentiates antituberculosis treatment via zinc poisoning in human macrophages and in a mouse model of infection. PLOS Biol. 22, e3002259 (2024).

67. Helaine, S., Conlon, B. P., Davis, K. M. & Russell, D. G. Host stress drives tolerance and persistence: The bane of anti-microbial therapeutics. Cell Host Microbe 32, 852–862 (2024).

68. Giraud-Gatineau, A. et al. The antibiotic bedaquiline activates host macrophage innate immune resistance to bacterial infection. eLife 9, e55692 (2020).

69. Coulson, G. B. et al. Targeting Mycobacterium tuberculosis Sensitivity to Thiol Stress at Acidic pH Kills the Bacterium and Potentiates Antibiotics. Cell Chem. Biol. 24, 993–1004.e4 (2017).

70. Dechow, S. J., Coulson, G. B., Wilson, M. W., Larsen, S. D. & Abramovitch, R. B. AC2P20 selectively kills Mycobacterium tuberculosis at acidic pH by depleting free thiols. RSC Adv. 11, 20089–20100 (2021).

71. Thiede, J. M. et al. Pyrazinamide Susceptibility Is Driven by Activation of the SigE-Dependent Cell Envelope Stress Response in Mycobacterium tuberculosis. mBio 13, e0043921 (2021).

72. Theriault, M. E. et al. Iron limitation in M. tuberculosis has broad impact on central carbon metabolism. *Commun*. Biol. 5, 1–11 (2022).

73. Manina, G., Dhar, N. & McKinney, J. D. Stress and Host Immunity Amplify Mycobacterium tuberculosis Phenotypic Heterogeneity and Induce Nongrowing Metabolically Active Forms. Cell Host Microbe 17, 32–46 (2015).

74. Liu, Y. et al. Immune activation of the host cell induces drug tolerance in Mycobacterium tuberculosis both in vitro and in vivo. J. Exp. Med. 213, 809–825 (2016).

75. Blondiaux, N. et al. Reversion of antibiotic resistance in Mycobacterium tuberculosis by spiroisoxazoline SMARt-420. Science 355, 1206–1211 (2017).

76. Kokoczka, R., Schuessler, D. L., Early, J. V. & Parish, T. Mycobacterium tuberculosis Rv0560c is not essential for growth in vitro or in macrophages. Tuberc. Edinb. Scotl. 102, 3–7 (2017).

77. Melief, E., Bonnett, S. A., Zuniga, E. S. & Parish, T. Activation of 2,4-Diaminoquinazoline in Mycobacterium tuberculosis by Rv3161c, a Putative Dioxygenase. Antimicrob. Agents Chemother. 63, e01505–18 (2019).

78. Gamngoen, R., Putim, C., Salee, P., Phunpae, P. & Butr-Indr, B. A comparison of Rv0559c and Rv0560c expression in drug-resistant Mycobacterium tuberculosis in response to first-line antituberculosis drugs. Tuberc. Edinb. Scotl. 108, 64–69 (2018).

79. Singh, A. et al. Requirement of the mymA operon for appropriate cell wall ultrastructure and persistence of Mycobacterium tuberculosis in the spleens of guinea pigs. J. Bacteriol. 187, 4173–4186 (2005).

80. Singh, S., Goswami, N., Tyagi, A. K. & Khare, G. Unraveling the role of the transcriptional regulator VirS in low pH-induced responses of Mycobacterium tuberculosis and identification of VirS inhibitors. J. Biol. Chem. 294, 10055–10075 (2019).

81. Vandal, O. H. et al. Acid-susceptible mutants of Mycobacterium tuberculosis share hypersusceptibility to cell wall and oxidative stress and to the host environment. J. Bacteriol. 191, 625–631 (2009).

82. Baker, J. J., Johnson, B. K. & Abramovitch, R. B. Slow growth of Mycobacterium tuberculosis at acidic pH is regulated by phoPR and host-associated carbon sources. Mol. Microbiol. 94, 56–69 (2014).

83. Prisic, S. et al. Extensive phosphorylation with overlapping specificity by Mycobacterium tuberculosis serine/threonine protein kinases. Proc. Natl. Acad. Sci. U. S. A. 107, 7521– 7526 (2010).

84. Gerrick, E. R. et al. Small RNA profiling in Mycobacterium tuberculosis identifies MrsI as necessary for an anticipatory iron sparing response. Proc. Natl. Acad. Sci. U. S. A. 115, 6464–6469 (2018).

85. Tan, S., Sukumar, N., Abramovitch, R. B., Parish, T. & Russell, D. G. Mycobacterium tuberculosis responds to chloride and pH as synergistic cues to the immune status of its host cell. PLoS Pathog. 9, e1003282 (2013).

86. Cortes, T. et al. Delayed effects of transcriptional responses in Mycobacterium tuberculosis exposed to nitric oxide suggest other mechanisms involved in survival. Sci. Rep. 7, 8208 (2017).

87. Chionh, Y. H. et al. tRNA-mediated codon-biased translation in mycobacterial hypoxic persistence. Nat. Commun. 7, 13302 (2016).

88. Arang, N. et al. Identifying host regulators and inhibitors of liver stage malaria infection using kinase activity profiles. Nat. Commun. 8, 1232 (2017).

89. Gujral, T. S., Peshkin, L. & Kirschner, M. W. Exploiting polypharmacology for drug target deconvolution. Proc. Natl. Acad. Sci. U. S. A. 111, 5048–5053 (2014).

90. Dankwa, S. et al. Exploiting polypharmacology to dissect host kinases and kinase inhibitors that modulate endothelial barrier integrity. Cell Chem. Biol. 28, 1679–1692.e4 (2021).

91. Anastassiadis, T., Deacon, S. W., Devarajan, K., Ma, H. & Peterson, J. R. Comprehensive assay of kinase catalytic activity reveals features of kinase inhibitor selectivity. Nat. Biotechnol. (2011) doi:10.1038/nbt.2017.

92. Klaeger, S. et al. The target landscape of clinical kinase drugs. Science 358, eaan4368 (2017).

93. Davis, M. I. et al. Comprehensive analysis of kinase inhibitor selectivity. Nat. Biotechnol. 29, 1046–1051 (2011).

94. Hie, B., Bryson, B. D. & Berger, B. Leveraging Uncertainty in Machine Learning Accelerates Biological Discovery and Design. Cell Syst. 11, 461–477.e9 (2020).

95. Hatzios, S. K., et al. Osmosensory signaling in Mycobacterium tuberculosis mediated by a eukaryotic-like Ser/Thr protein kinase. Proc. Natl. Acad. Sci. 110, E5069–E5077 (2013).

96. Canton, J., Khezri, R., Glogauer, M. & Grinstein, S. Contrasting phagosome pH regulation and maturation in human M1 and M2 macrophages. Mol. Biol. Cell (2014) doi:10.1091/mbc.E14-05-0967.

97. Teplova, V. V., Tonshin, A. A., Grigoriev, P. A., Saris, N.-E. L. & Salkinoja-Salonen, M. S. Bafilomycin A1 is a potassium ionophore that impairs mitochondrial functions. J. Bioenerg. Biomembr. 39, 321–329 (2007).

98. Rybicka, J. M., Balce, D. R., Khan, M. F., Krohn, R. M. & Yates, R. M. NADPH oxidase activity controls phagosomal proteolysis in macrophages through modulation of the lumenal redox environment of phagosomes. Proc. Natl. Acad. Sci. U. S. A. 107, 10496–10501 (2010).

99. Burger, H. et al. Lysosomal Sequestration Determines Intracellular Imatinib Levels. Mol. Pharmacol. 88, 477–487 (2015).

100. Rybniker, J. et al. Anticytolytic screen identifies inhibitors of mycobacterial virulence protein secretion. Cell Host Microbe 16, 538–548 (2014).

101. Culviner, P. H. et al. Evolution of Mycobacterium tuberculosis transcription regulation is associated with increased transmission and drug resistance. bioRxiv 2025.05.01.651750 (2025) doi:10.1101/2025.05.01.651750.

102. Parekh, S., Ziegenhain, C., Vieth, B., Enard, W. & Hellmann, I. zUMIs - A fast and flexible pipeline to process RNA sequencing data with UMIs. GigaScience 7, giy059 (2018).

103. Love, M. I., Huber, W. & Anders, S. Moderated estimation of fold change and dispersion for RNA-seq data with DESeq2. Genome Biol. 15, 550 (2014).

104. Golby, P. et al. Characterization of two in vivo-expressed methyltransferases of the Mycobacterium tuberculosis complex: antigenicity and genetic regulation. Microbiology 154, 1059–1067 (2008).

